# Elevated glycolytic metabolism of monocytes limits the generation of HIF-1α-driven migratory dendritic cells in tuberculosis

**DOI:** 10.1101/2023.04.03.535400

**Authors:** Mariano Maio, Joaquina Barros, Marine Joly, Zoi Vahlas, José Luis Marín Franco, Melanie Genoula, Sarah Monard, María Belén Vecchione, Federico Fuentes, Virginia Gonzalez Polo, María Florencia Quiroga, Mónica Vermeulen, Thien-Phong Vu Manh, Rafael J Argüello, Sandra Inwentarz, Rosa Musella, Lorena Ciallella, Pablo González Montaner, Domingo Palmero, Geanncarlo Lugo Villarino, María del Carmen Sasiain, Olivier Neyrolles, Christel Verollet, Luciana Balboa

**Affiliations:** Instituto de Medicina Experimental (IMEX)-CONICET, Academia Nacional de Medicina, Buenos Aires, Argentina; International Associated Laboratory (LIA) CNRS IM-TB/HIV (1167), Buenos Aires, Argentina / International Research Project Toulouse, France; Instituto de Investigaciones Biomédicas en Retrovirus y Sida (INBIRS), Consejo Nacional de Investigaciones Científicas y Técnicas (CONICET) - Universidad de Buenos Aires. Buenos Aires, Argentina; Institut de Pharmacologie et de Biologie Structurale, Université de Toulouse, CNRS, UPS, Toulouse, France; Aix Marseille University, CNRS, INSERM, CIML, Centre d’Immunologie de Marseille-Luminy, Marseille, France; Instituto Prof. Dr. Raúl Vaccarezza and Hospital de Infecciosas Dr. F.J. Muñiz, Buenos Aires, Argentina

## Abstract

During tuberculosis, migration of dendritic cells (DCs) from the site of infection to the draining lymph nodes is known to be impaired, hindering the rapid development of protective T-cell mediated immunity. However, the mechanisms involved in the delayed migration of DCs during tuberculosis (TB) are still poorly defined. Here, we found that infection of DCs with *Mycobacterium tuberculosis* (Mtb) triggers HIF-1α-mediated aerobic glycolysis in a TLR2-dependent manner, and that this metabolic profile is essential for DC migration. In particular, the lactate dehydrogenase (LDH) inhibitor oxamate and the HIF-1α inhibitor PX-478 abrogated Mtb-induced DC migration *in vitro* to the lymphoid tissue-specific chemokine CCL21, and *in vivo* to lymph nodes in mice. Strikingly, we found that although monocytes from TB patients are inherently biased toward glycolysis metabolism, they differentiate into poorly glycolytic and poorly migratory DCs, compared with healthy subjects. Taken together, these data suggest that because of their preexisting glycolytic state, circulating monocytes from TB patients are refractory to differentiation into migratory DCs, which may explain the delayed migration of these cells during the disease and opens avenues for host-directed therapies for TB.

**Graphical Abstract:** 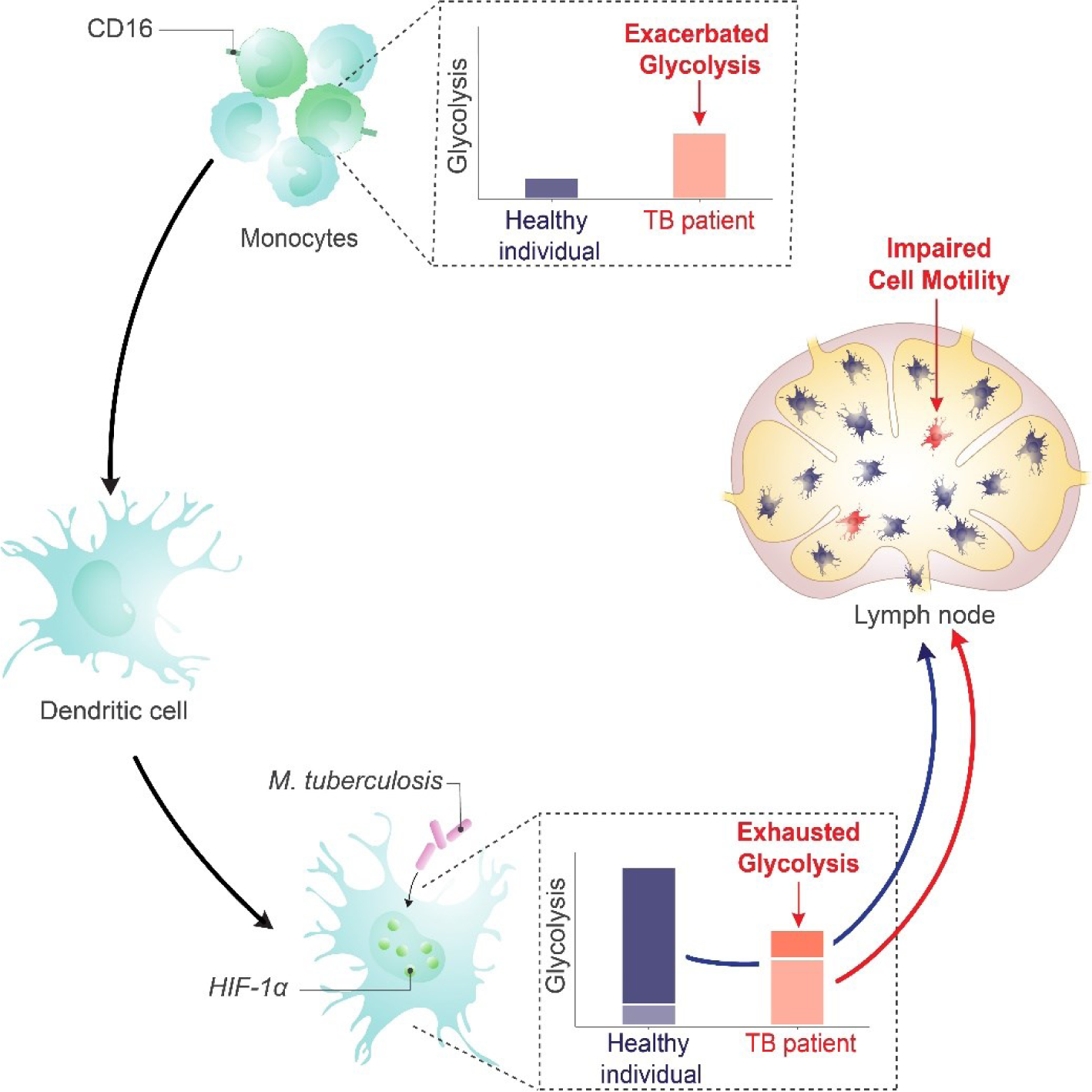

## Introduction

Tuberculosis (TB) remains a major global health problem, responsible for approximately 1.6 million deaths annually. The causative agent of TB, *Mycobacterium tuberculosis* (Mtb), is a highly successful pathogen that has evolved several strategies to weaken the host immune response. Although reliable immune correlates of protective immunity against Mtb are still not well-defined, it is widely accepted that Th1 cells contribute to protection by secreting IFN-γ and promoting antimycobacterial activity in macrophages^1^. Importantly, the induction of a strong Th1 immune response relies on the generation of immunogenic dendritic cells (DCs) with strong migratory properties^2–5^. Mtb has been shown to interfere with several DC functions, thus impairing the induction and development of adaptive immunity^6–8^. For instance, we and others previously reported that Mtb-exposed DCs have low capacity for mycobacterial antigen presentation and stimulation of Mtb-specific CD4^+^ T cells^9–13^. Additionally, Mtb-infected DCs were reported to have an impaired ability to migrate to lymph nodes *in vitro*^14,15^ and *in vivo* in murine models^3,5^; however, the underlying molecular mechanisms of these phenotypes and their relevance to the migratory activity of monocyte-derived DCs in TB patients remain unknown.

Rapid, directed migration of DCs towards secondary lymphoid organs requires essential changes at the cellular and molecular levels^16^. Relatedly, the metabolic state of DCs is complex and varies according to cell origin, differentiation and maturation states, as well as local microenvironment, among other factors^17–20^. Studies have reported that upon pathogen sensing, the transcription factor hypoxia-inducible factor-1α (HIF-1α) increases glycolysis, which promotes immunogenic functions of DCs, such as IL-12 production, costimulatory marker expression^21^, and cell migration^22–24^. By contrast, it was shown that HIF-1α represses the proinflammatory output of LPS-stimulated DCs and can inhibit DC-induced T-cell responses in other settings^25^. To reconcile these disparate roles for HIF-1α, it has been proposed that the impact of metabolic pathway activation on DC functions varies among DC subsets^18^. To this point, most prior studies have been conducted using murine conventional DCs and plasmacytoid DCs^19^. Recently, with the implementation of high-dimensional techniques, it was demonstrated that distinct metabolic wiring is associated with individual differentiation and maturation stages of DCs^26^, highlighting the importance of defining the metabolic profile of specific subsets of DCs under physiological or pathological conditions^20^. Given the key role of DCs in the host response to TB, it is thus crucial to investigate DC metabolism in the context of Mtb infection^27^.

We previously demonstrated that the TB-associated microenvironment, as conferred by the acellular fraction of TB patient pleural effusions, inhibits HIF-1α activity leading to a reduction in glycolytic and microbicidal phenotypes in macrophages^28^. Moreover, activation of HIF-1α enhances Mtb control at early times post-infection in mouse models^29^, and this effect was associated with a metabolic switch of alveolar macrophages towards an M1-like profile^28^. Given that HIF-1α activation promotes protection at early stages of Mtb infection and given its role as a key regulator of DC migration and inflammation^30^, we hypothesized that HIF-1α could affect the functionality of DCs in regulating the initiation and orchestration of the adaptive immune response to Mtb, a process known to be delayed upon Mtb infection^5,6^. Here, we show that HIF-1α-mediated glycolysis promotes DC activation and migration in the context of TB. Importantly, we report active glycolysis in monocytes from TB patients, which leads to poor glycolytic induction and migratory capacities of monocyte-derived DCs.

## Results

### Mtb impacts metabolism in human monocyte-derived DCs

To determine the impact of Mtb on the metabolism of human monocyte-derived DCs (Mo-DCs), we assessed metabolic parameters associated with glycolysis and mitochondrial changes upon Mtb stimulation or infection. Cells undergoing aerobic glycolysis are characterized by increased consumption of glucose and the production and release of lactate. We measured lactate release and glucose consumption in Mo-DCs stimulated for 24 h with equivalent doses of either irradiated (iMtb) or viable Mtb. DCs treated with either iMtb or viable Mtb released increased levels of lactate and consumed more glucose than untreated DCs (**Figure 1A-B**). Consistently, both iMtb treatment and Mtb infection resulted in an increase in expression of the key glycolysis-activating regulator HIF-1α at both mRNA and protein levels (**Figure 1C-D**). Expression of the gene encoding the glycolytic enzyme lactate dehydrogenase A (*LDHA*), which catalyzes the conversion of lactate to pyruvate, was also increased in iMtb-treated or Mtb-infected DCs (**Figure 1E**). In agreement with their enhanced glycolysis profile, DCs stimulated with iMtb or infected with viable Mtb had increased expression of the glucose transporter GLUT1 (SLC2A1)^31^ (**Figure 1F**). Of note, *LDHA* and *GLUT1* are HIF-1α target genes, and their upregulation correlated with the increase in HIF-1α expression upon Mtb stimulation. To assess changes in the mitochondria, we measured mitochondrial mass and morphology. We found a higher mitochondrial mass as well as larger individual mitochondria in iMtb-stimulated DCs compared to untreated DCs (**Figure 1G-H**). In contrast to the findings obtained upon iMtb stimulation, Mtb-infected DCs displayed a reduction in their mitochondrial mass (**Figure 1G**). This result indicates that although both Mtb-infected and irradiated Mtb-exposed DCs show a clear increase in their glycolytic activity, divergent responses are observed in terms of mitochondrial mass. Therefore, our data indicate that Mtb impacts the metabolism of Mo-DCs, leading to mitochondrial changes and triggering glycolysis-associated parameters.

**Figure 1.**
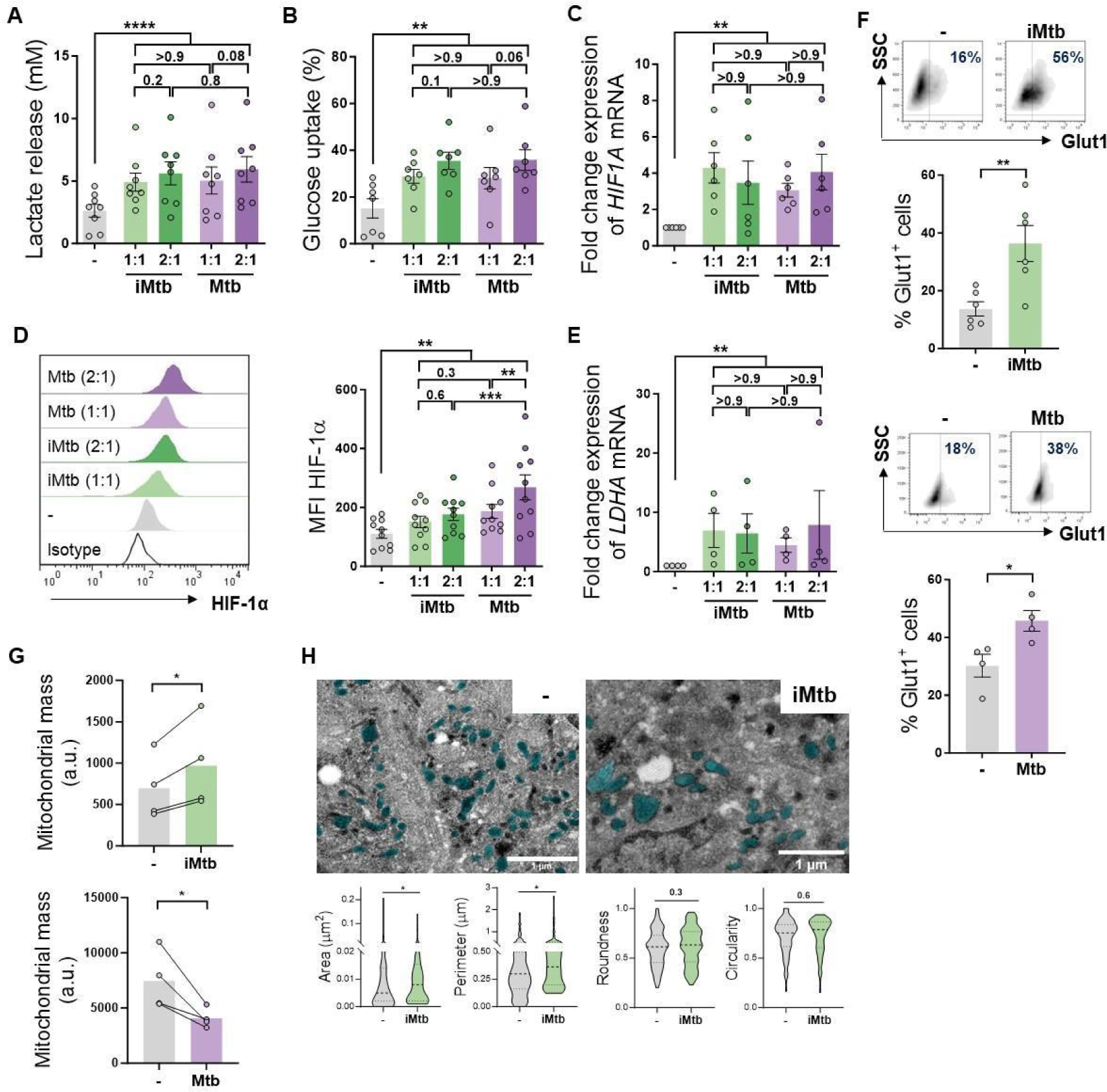
Mtb rewires the metabolic network of monocyte-derived DCs (Mo-DCs). Mo-DCs were stimulated with viable or irradiated Mtb (iMtb) at two MOI (1 or 2 Mtb per DC) for 24 h. Glycolysis was measured as: **(A)** Lactate release in culture supernatants (N=8); **(B)** Glucose uptake measured in culture supernatants (N=7); **(C)** Relative expression of *HIF-1α* mRNA normalized to *EeF1A1* control gene (N=6). **(D)** Representative histograms of the mean fluorescence intensity (MFI) of HIF-1α as measured by flow cytometry. Quantification shown in graph to the right (N=10). **(E)** Relative expression of lactate dehydrogenase A (*LDHA*) mRNA normalized to *EeF1A1* control gene (N=4). **(F)** FACS plots show the percentage of Glut1^+^ cells with and without iMtb stimulation or infected with viable Mtb in a representative experiment. Quantification of Glut1+ cells plotted below (N=4-6). **(G)** MFI of Mitospy probe as a measurement of mitochondrial mass for Mo-DCs treated (or not) with iMtb (upper panel) or infected with viable Mtb (lower panel). The data are represented as scatter plots with each circle representing a single individual, means ± SEM are shown (N=4). **(H)** Representative electron microscopy micrographs of control and iMtb-stimulated DCs showing mitochondria colored in cyan (left panels) and quantified morphometric analysis (right panels) (N=4). Statistical significance was assessed in **(A-E)** using 2-way ANOVA followed by Tukey’s multiple comparisons test (∗p < 0.05; ∗∗p < 0.01; ∗∗∗∗p < 0.0001), and in **(F-H)** using paired t test (∗p < 0.05) for iMtb versus controls. All values are expressed as means ± SEM.

### Mtb exposure shifts DCs to a glycolytic profile over oxidative phosphorylation

To further characterize the metabolic profile of DCs upon iMtb-stimulation or Mtb-infection, we next evaluated the metabolism of DCs at single-cell level using the SCENITH technology^32^. This method is based on a decrease in ATP levels that is tightly coupled with a decrease in protein synthesis and displays similar kinetics^32^. By treating the cells with glucose or mitochondrial respiration inhibitors, and measuring their impact on protein synthesis by puromycin incorporation via flow cytometry, glucose and mitochondrial dependences can be quantified. Two additional derived parameters such as “glycolytic capacity” and “fatty acid and amino acid oxidation (FAO & AAO) capacity” were also calculated. SCENITH technology revealed a lower reliance on oxidative phosphorylation (OXPHOS) in parallel with an increase in the glycolytic capacity of iMtb-stimulated (**Figure 2A-B**), Mtb-infected DCs and even bystander DCs (those cells that are not directly infected but stand nearby) (**Figure 2C-D**). Since bystander DCs are not in direct association with Mtb (Mtb-RFP-DCs), soluble mediators induced in response to infection may be sufficient to trigger glycolysis even in uninfected cells. No differences were observed for glucose dependence and FAO & AAO capacity (**Figure 2A-D**). Additionally, we found no changes between the FAO dependency in Mtb-stimulated DCs in comparison to control cells when the FAO inhibitor (etomoxir) was used (**Figure S1A**). For the case of iMtb-stimulated DCs, we also assessed the intracellular rates of glycolytic and mitochondrial ATP production using Seahorse technology. Bioenergetic profiles revealed that iMtb increased the rate of protons extruded over time, or proton efflux rate (PER), as well as the basal oxygen consumption rate (OCR) in Mo-DCs (**Figure 2E**). The measurements of basal extracellular acidification rate (ECAR) and OCR were used to calculate ATP production rate from glycolysis (GlycoATP) and mitochondrial OXPHOS (MitoATP). The ATP production rates from both glycolysis and mitochondrial respiration were augmented upon iMtb-stimulation (**Figure 2F**). Similar to SCENITH results, the relative contribution of GlycoATP to overall ATP production was increased, while MitoATP contribution was decreased in iMtb-treated cells compared to untreated cells (**Figure 2G**). These results confirmed the change in DC metabolism induced by Mtb, with an increase in the relative glycolytic contribution to overall metabolism at the expense of the OXPHOS pathway. Together, metabolic profiling indicates that a metabolic switch toward aerobic glycolysis occurs in Mo-DCs exposed to Mtb.

**Figure 2.**
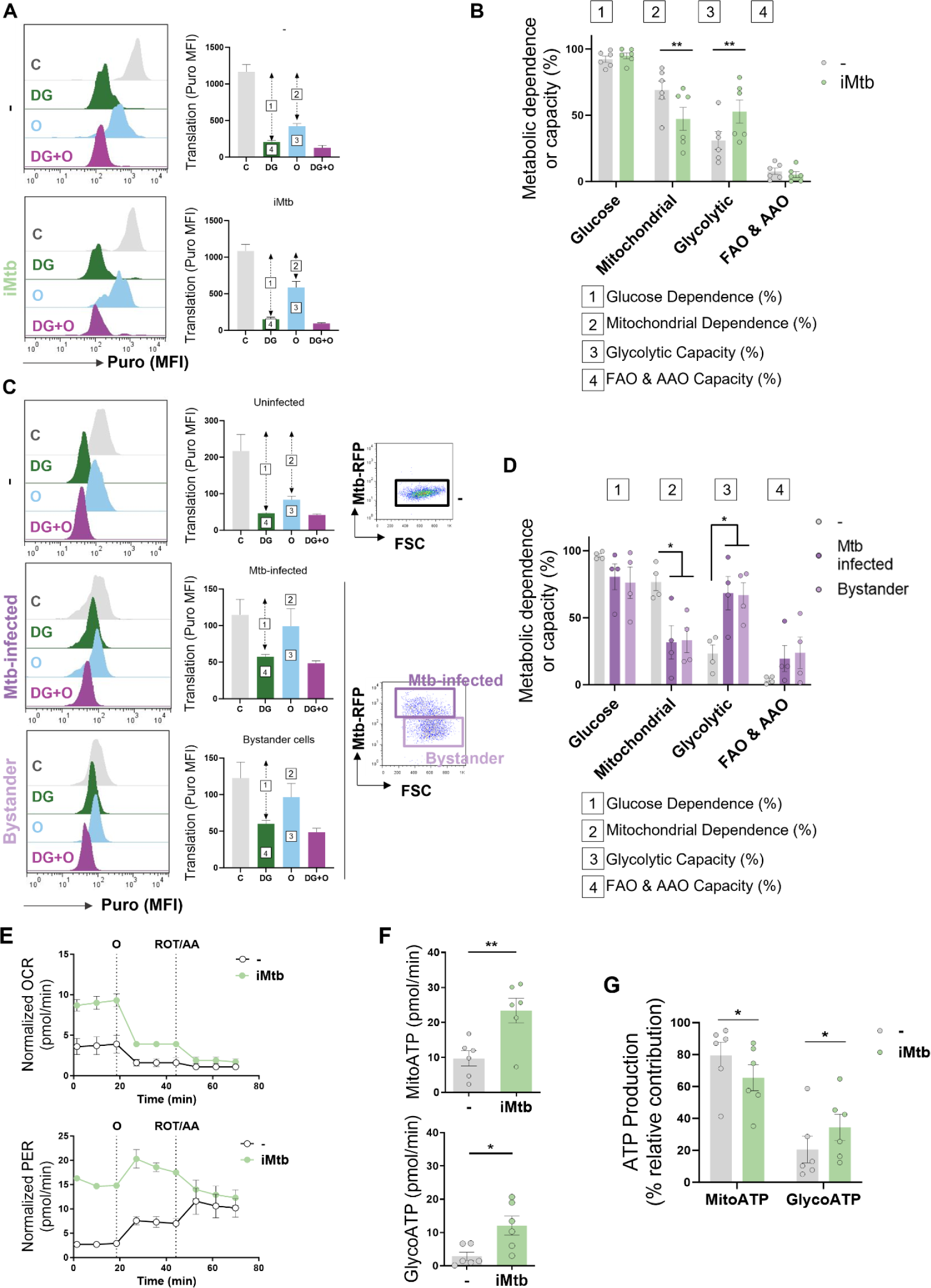
Mtb skews DC metabolism toward glycolysis. Mo-DCs were stimulated with irradiated Mtb (iMtb) or infected with Mtb expressing Red Fluorescent Protein (Mtb-RFP, panel C). **(A)** Representative histograms showing the translation level after puromycin (Puro) incorporation and staining with a monoclonal anti-Puro (anti-Puro MFI) in response to inhibitor treatment (C, Control; DG, 2-Deoxy-D-Glucose; Oligomycin, O; or combination treatment, DG+O). The bar plots show the values of the anti-Puro MFI from 6 donors. Arrows and numbers inside boxes denote the differences between the MFI of puro in the different treatments that are used to calculate the glucose dependence (1) and fatty acids and amino acids oxidation (FAO & AAO) capacity (4); and the mitochondrial dependency (2) and glycolytic capacity (3). **(B)** Relative contributions of glycolytic and FAO & AAO capacities and glucose and mitochondrial dependences to overall DC metabolism analyzed with SCENITH (N=6)**. (C-D)** DCs were infected with Mtb-RFP for 24 h, thereafter the metabolic profile was evaluated by SCENITH. **(C)** Representative histograms showing the translation level after Puro incorporation are shown for uninfected, Mtb-infected and bystander DCs (those cells that are not infected directly but rather stand nearby). The bar plots show the values of the anti-Puro MFI from 4 donors. Right panel shows representative plots showing the gating strategy to distinguish the populations within Mtb-infected cultures, which includes RFP^+^ (Mtb-infected DCs) and RFP^-^ (bystander DCs) cells. **(D)** Relative contributions of glycolytic and FAO & AAO capacities and glucose and mitochondrial dependences to DC metabolism (N=4). **(E)** Kinetic profile of proton efflux rate (PER; lower panel) and oxygen consumption rate (OCR; upper panel) measurements in control and iMtb-stimulated DCs in response to inhibitor treatments (Oligomycin, O; ROT/AA, Rotenone/Antimycin A), obtained using an Agilent Seahorse XFe24 Analyzer. PER and OCR measurements were normalized to the area covered by cells. **(F)** ATP production rate from mitochondrial oxidative phosphorylation (MitoATP) and glycolysis (glycoATP). MitoATP production rate and glycoATP production rate were calculated from OCR and ECAR measurements in control and iMtb-stimulated DCs (N=6). **(G)** Percentages of MitoATP and GlycoATP relative to overall ATP production (N=6). Statistics in **(B, F-G)** are from paired t test (∗p < 0.05; ∗∗p < 0.01) for iMtb versus controls. Statistics in **(D)** are 2-way ANOVA followed by Tukey’s multiple comparisons test (∗p < 0.05) as depicted by lines. The data are represented as scatter plots with each circle representing a single individual, means ± SEM are shown.

### Mtb triggers the glycolytic pathway through TLR2 ligation

Since Mtb is sensed by Toll-like receptors (TLR)-2 and −4^33^, we investigated the contribution of these receptors to glycolysis activation in Mo-DCs upon Mtb stimulation. Using specific neutralizing antibodies for these receptors, we found that TLR2 ligation, but not that of TLR4, was required to trigger the glycolytic pathway, as reflected by a decrease in lactate release, glucose consumption and HIF-1α expression in iMtb-stimulated DCs treated with an anti-TLR2 antibody (**Figure 3A-C**). As a control, and as expected given the reliance on TLR4 for LPS sensing^34^, lactate release and glucose consumption were abolished in LPS-stimulated DCs in the presence of neutralizing antibodies against TLR4 but not TLR2 (**Figure S2A-B**). Moreover, blockade of TLR2 also diminished glycolytic ATP production in iMtb-stimulated DCs without altering OXPHOS-associated ATP production (**Figure 3D**) or the size and morphology of mitochondria (**Figure S2C**), suggesting that TLR2 engagement by iMtb is required for the induction of glycolysis but not mitochondrial respiration. Interestingly, TLR2 ligation was also necessary for lactate release and HIF-1α up-regulation triggered by viable Mtb, (**Figure S2D-E**). To further confirm the involvement of TLR2 in the induction of glycolysis, we tested the effect of synthetic (Pam_3_CSK_4_) or mycobacterial (peptidoglycans, PTG) TLR2 agonists^35,36^ and found that both ligands induced lactate release and glucose consumption in DCs (**Figure 3E-F**), without affecting cell viability (**Figure S2F**). Thus, our data indicate that Mtb induces glycolysis in Mo-DCs through TLR2 engagement.

**Figure 3.**
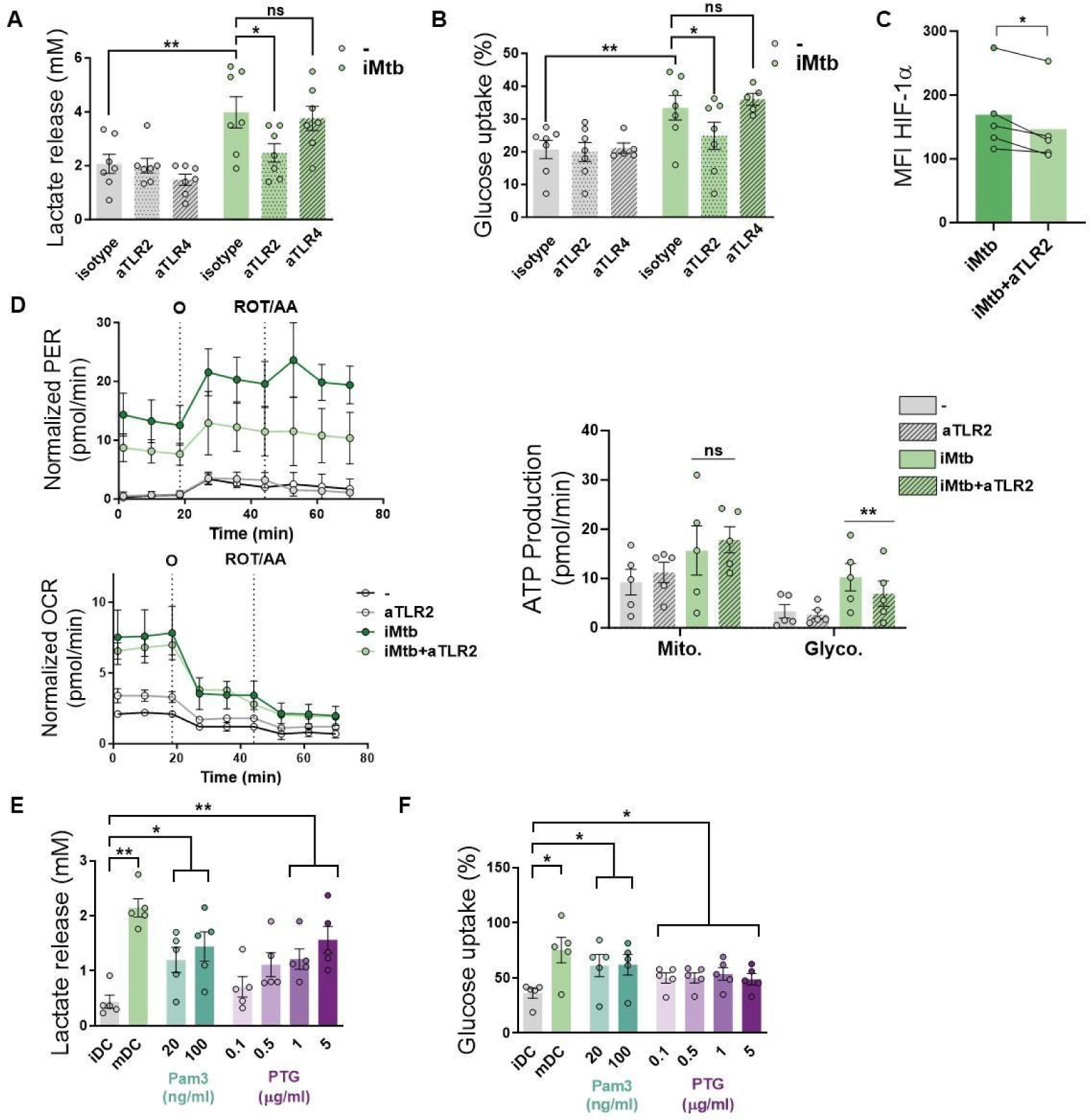
Mtb triggers glycolysis through TLR2 ligation in Mo-DCs. Mo-DCs were stimulated with irradiated Mtb (iMtb) in the presence of neutralizing antibodies against either TLR2 (aTLR2), TLR4 (aTLR4), or their respective isotype controls. **(A)** Lactate release as measured in supernatant (N=7). **(B)** Glucose uptake as measured in supernatant (N=7). **(C)** Mean fluorescence intensity (MFI) of HIF-1α as measured by flow cytometry (N=4). **(D)** Kinetic profile of proton efflux rate (PER) and oxygen consumption rate (OCR) measurements (left panels). Metabolic flux analysis showing quantification of mitochondrial ATP production and glycolytic ATP production (right panel) (N=5). **(E-F)** Mo-DCs were stimulated with Pam3Cys or Mtb peptidoglycan (PTG) at the indicated concentrations. **(E)** Lactate release as measured in supernatant (N=5). **(F)** Glucose uptake as measured in supernatant (N=5). Statistics in **(A-B, E-F)** are 2-way ANOVA followed by Tukey’s multiple comparisons test (∗p < 0.05; ∗∗p < 0.01; ∗∗∗∗p < 0.0001). Statistics in **(C-D)** are from paired t test (∗p < 0.05) for iMtb versus controls. The data are represented as scatter plots with each circle representing a single individual, means ± SEM are shown.

### HIF-1α is required for DC maturation upon iMtb stimulation but not for CD4^+^ T lymphocyte polarization

To determine the impact of glycolysis on DC maturation and the capacity to activate T cells, we inhibited HIF-1α activity in iMtb-stimulated DCs employing two HIF-1α inhibitors that display different mechanisms of action. The first is PX-478 (PX) that lowers HIF-1α levels by inhibiting HIF-1α deubiquitination, decreases HIF-1α mRNA expression, and reduces HIF-1α translation^37^; the second one is echinomycin (Ech), which inhibits the binding of HIF-1α to the hypoxia response element thereby blocking HIF-1α DNA binding capability^38,39^. Treatment with either HIF-1α inhibitor PX or Ech diminished lactate release and glucose consumption in iMtb-stimulated DCs without affecting cell viability at the indicated concentration (**Figure S3A-F**). HIF-1α inhibition by PX significantly abolished ATP production associated with glycolysis without affecting absolute levels of OXPHOS-derived ATP production in iMtb-stimulated DCs (**Figure 4A and Figure S3G**). In line with these results, the glycolytic capacity was reduced in iMtb-stimulated DCs treated with Ech (**Figure S3H**). Although HIF-1α inhibitors did not affect the uptake of iMtb by DCs (**Figure S3I**), we observed a reduction in the expression of activation markers CD83 and CD86, but not in the inhibitory molecule PD-L1, upon treatment with PX (**Figure 4B**) or Ech (**Figure S4**). We then measured cytokine production in iMtb-stimulated DCs after HIF-1α inhibition and noted a reduction in TNF-α and an increase in IL-10 production by PX-treated cells (**Figure 4C**). To assess the capacity of DCs to activate T cells in response to mycobacterial antigens, we co-cultured DCs and autologous CD4^+^ T cells from PPD^+^ donors in the presence or absence of HIF-1α inhibitors and measured the overall IFN-γ and IL-17 production in culture supernatants as well as cell surface expression of T cell markers. We found no significant differences in the activation profile of autologous CD4^+^ T cells in coculture with iMtb-stimulated DCs treated or not with PX (**Figure 4D-E and Figure S5**). We conclude that while HIF-1α is important for the maturation of iMtb-stimulated Mo-DCs, it does not influence their capacity to activate CD4^+^ T cells, at least *in vitro*.

**Figure 4.**
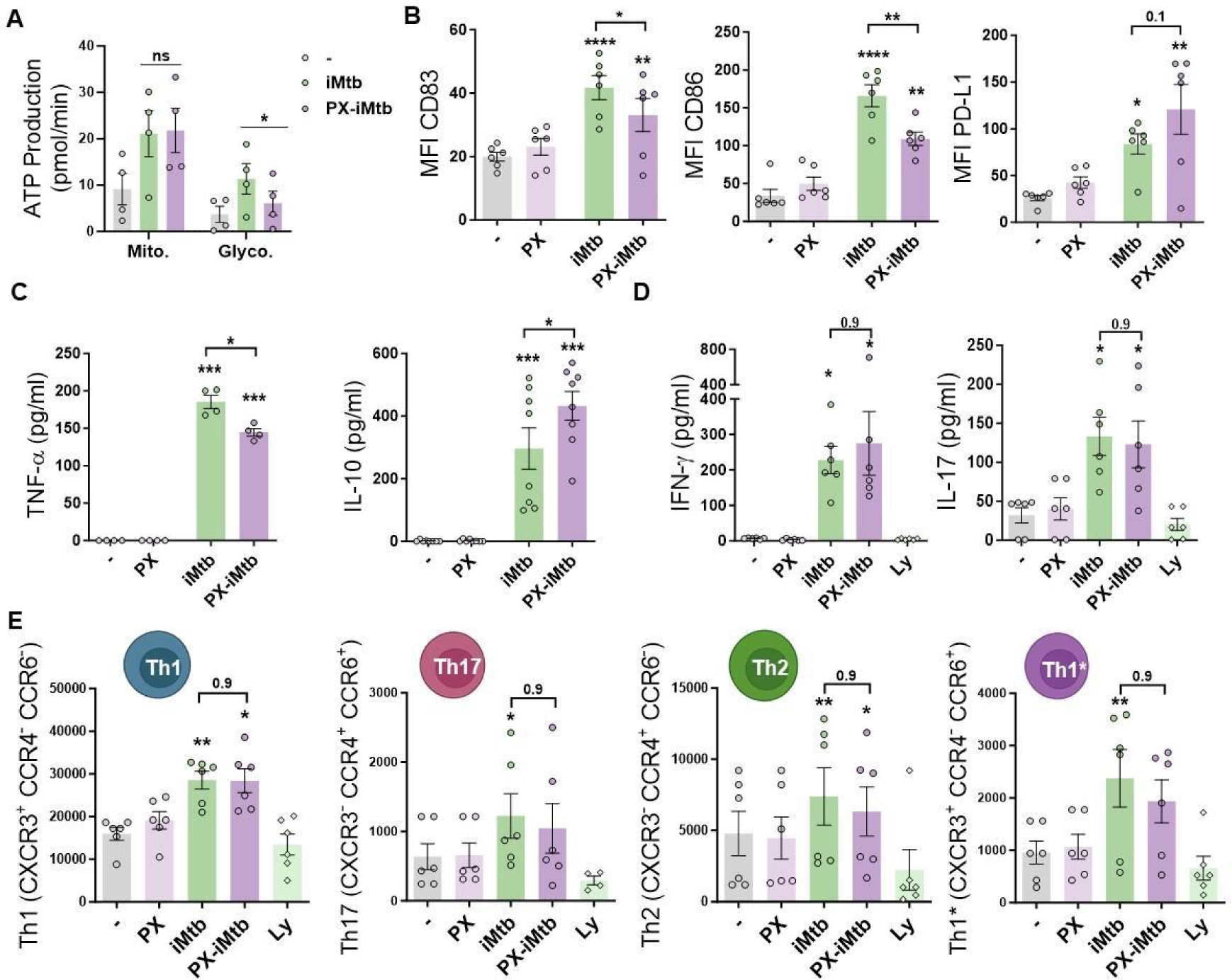
HIF-1α is required for DC maturation upon iMtb stimulation but not for CD4^+^ T lymphocyte polarization. **(A-C)** Mo-DCs were stimulated with irradiated Mtb (iMtb) in the presence or absence of the HIF-1α inhibitor PX-478 (PX). **(A)** Metabolic flux analysis showing quantification of mitochondrial ATP production and glycolytic ATP production, as in Figure 2C (N=4). **(B)** Mean fluorescence intensity (MFI) of CD83, CD86 and PD-L1 as measured by flow cytometry (N=6). **(C)** TNF-α and IL-10 production by Mo-DCs measured by ELISA (N=4-8). **(D-E)** Monocytes from PPD^+^ healthy donors were differentiated towards DCs, challenged or not with iMtb in the presence or absence of PX for 24 h, washed, and co-cultured with autologous CD4^+^ T cells for 5 days. **(D)** Extracellular secretion of IFN-γ and IL-17 as measured by ELISA (N=6). **(E)** Absolute abundance of Th1, Th17, Th2 and Th1/Th17 CD4^+^ T cells after coculture with DCs (N=6). When indicated lymphocytes without DCs were cultured (Ly). Statistical significance based on 2-way ANOVA followed by Tukey’s multiple comparison test (∗p < 0.05; ∗∗p < 0.01). The data are represented as scatter plots with each circle representing a single individual, means ± SEM are shown.

### HIF-1α-mediated glycolysis triggers the motility of DCs upon iMtb stimulation

Since DC migration to lymph nodes is essential to initiate an adaptive immune response and glycolytic activity has been reported to control DC migration upon stimulation^22,23^, we evaluated the migratory properties of iMtb-stimulated DCs in the presence of inhibitors of HIF-1α and LDH which catalyzes the interconversion of pyruvate and lactate. First, we confirmed that PX and oxamate (OX), a well-established LDH inhibitor, diminished the glycolytic activity of iMtb-stimulated human Mo-DCs, as demonstrated by reduced lactate release (**Figure S3A and 5A**). Next, using a transwell migration assay, we found that PX and OX treatment significantly diminished the chemotactic activity of iMtb-stimulated human Mo-DCs in response to CCL21 (**Figure 5B**), a CCR7 ligand responsible for the migration of DCs into lymphoid organs. We also assessed the 3 dimensional (3D) migration capacity of iMtb-stimulated DCs through a collagen matrix in which DCs use an amoeboid migration mode^40^ and found that 3D migration was significantly impaired upon HIF-1α or glycolysis inhibition (**Figure 5C**). The role of glycolysis in the migration of iMtb-stimulated Mo-DCs was further confirmed using an additional LDHA inhibitor, GSK2837808A, which reduced both the release of lactate by iMtb-stimulated Mo-DCs and their migration in response to CCL21 (**Figure S6A-B**). Attenuation of cell migration through collagen induced by OX and PX was also confirmed in Mtb-infected DCs (**Figure 5D**). To further investigate the effects of glycolysis on cell migration, we turned to an *in vivo* model. Murine bone marrow-derived DCs (BMDCs) isolated and stimulated with iMtb in the presence or absence of PX or OX were labeled with CFSE and transferred into naïve mice (**Figure 5E**). Similar to human Mo-DCs, iMtb stimulation increased glycolysis in BMDCs, which was inhibited by PX and OX treatment *in vitro* (**Figure S6C**). Three hours after the transfer of BMDCs into recipient mice, nearby lymph nodes were collected for DC quantification (**Figure 5E**). A higher number of adoptively transferred DCs (CFSE-labeled CD11c^+^ cells) were detected in lymph nodes from mice that received iMtb-stimulated BMDCs compared to mice that received untreated BMDCs or iMtb-BMDCs treated with either PX or OX (**Figure 5F and Figure S6D**). Of note, we verified that CCR7 expression on iMtb-stimulated BMDCs was not affected by OX or PX treatment, so the effect could not be ascribed to downregulation of the chemokine receptor (**Figure S6E**). Therefore, we conclude that HIF-1α-mediated glycolysis is required for the successful migration of iMtb-stimulated DCs into lymph nodes.

**Figure 5.**
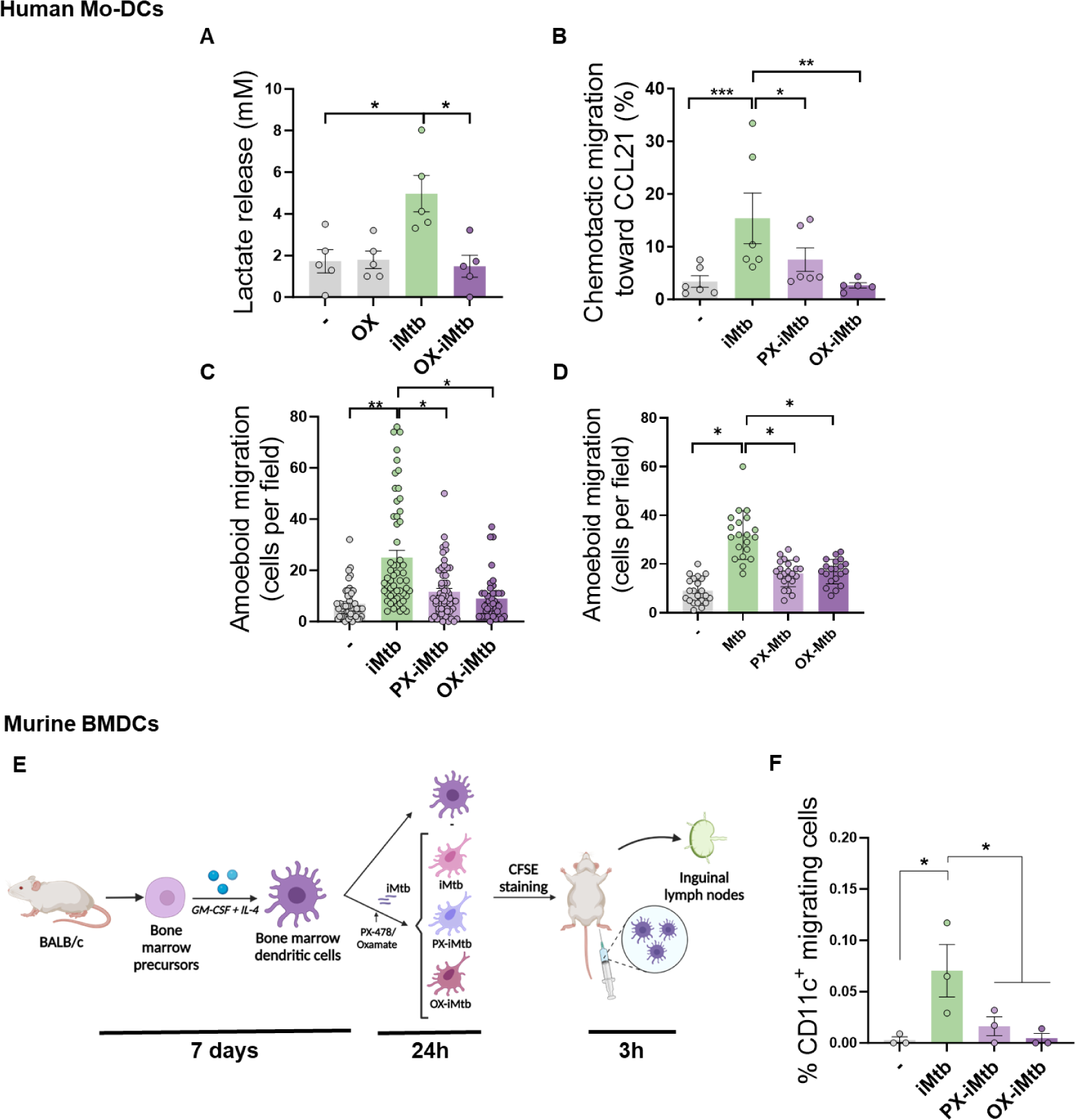
HIF1α-mediated-glycolysis is required to trigger migratory activity in iMtb-stimulated DCs. Mo-DCs were treated (or not) with HIF-1a inhibitor PX-478 (PX) or LDH inhibitor oxamate (OX) and stimulated with iMtb for 24 h. **(A)** Lactate release as measured in supernatants in DCs stimulated or not with iMtb in the presence of OX (N=5). **(B)** Percentage of migrated cells towards CCL21 relative to the number of initial cells per condition (N=6). **(C-D)** Three-dimensional amoeboid migration of DCs through a collagen matrix after 24 h. Cells within the matrix were fixed and stained with DAPI. Images of the membrane of each insert were taken and the percentage of cells per field were counted. **(C)** Mo-DCs stimulated with iMtb for 24 h (N=5). **(D)** Mo-DCs infected with Mtb for 24 h (N=4). The data are represented as scatter plots, where each circle represents a microphotograph sourced from either 5 (C) or 4 (D) independent donors, with each experiment typically including between five to ten microphotographs. **(E)** Representative schematic of the experimental setup for *in vivo* migration assays. **(F)** Percentages of migrating BMDCs (CFSE-labeled among CD11c^+^) recovered from inguinal lymph nodes (N=3). Statistical significance assessed by **(A-B)** ANOVA followed by Dunnett’s multiple comparisons test (∗p < 0.05; ∗∗p < 0.01); **(C-D)** Nested ANOVA followed by Dunnett’s multiple comparisons test (∗p < 0.05; ∗∗p < 0.01); **(F)** ANOVA followed by Holm-Sidak’s multiple comparisons test (∗p < 0.05).

### Stabilization of HIF-1α promotes migration of tolerogenic DCs and DCs derived from TB patient monocytes

Since DC differentiation is skewed, at least partially, towards a tolerogenic phenotype during TB^41–43^, we investigated whether tolerogenic DCs can be reprogrammed into immunogenic DCs by modulating their glycolytic pathway after iMtb stimulation. To this end, we generated tolerogenic Mo-DCs by adding dexamethasone (Dx) before stimulation with iMtb in the presence or absence of dimethyloxalylglycine (DMOG), which stabilizes the expression of HIF-1α. HIF-1α expression is tightly regulated by prolyl hydroxylase domain containing proteins which facilitate the recruitment of the von Hippel-Lindau (VHL) protein, leading to ubiquitination and degradation of HIF-1α by the proteasomes^44^. DMOG inhibits the prolyl hydroxylase domain-containing proteins. Acquisition of the tolerogenic phenotype was confirmed by the lack of upregulation of costimulatory markers CD83 and CD86, as well as by increased PD-L1 expression in iMtb-DCs treated with Dx compared to control iMtb-DCs (**Figure S7A**). Moreover, Dx-treated DCs did not exhibit an increase in lactate release, consumption of glucose or induction of HIF-1α expression in response to iMtb, showing a high consumption of levels of glucose under basal conditions (**Figure 6A-B**). Of note, HIF-1α stabilization using DMOG restored the HIF-1α expression and lactate production in response to iMtb in Dx-treated DCs and increased the consumption of glucose (**Figure 6A-B**). Activation of HIF-1α also improved 3D amoeboid migration, as well as 2D migration capacity of DCs towards CCL21 of iMtb-stimulated Dx-treated DCs (**Figure 6C-D and Figure S7B**). Confirming the relevance of these findings to human TB patients, we found that iMtb-stimulated Mo-DCs from TB patients were deficient in their capacity to migrate towards CCL21 (**Figure 6E**) and in glycolytic activity, compared to Mo-DCs from healthy subjects (**Figure 6F-G**). Strikingly, stabilizing HIF-1α expression using DMOG in Mo-DCs from TB patients restored their chemotactic activity in response to iMtb (**Figure 6H**). These data indicate that the impaired migratory capacity of iMtb-stimulated tolerogenic DCs or TB patient-derived DCs can be restored via HIF-1α stabilization; thus, glycolysis is critical for DC function during TB in both murine and human contexts.

**Figure 6.**
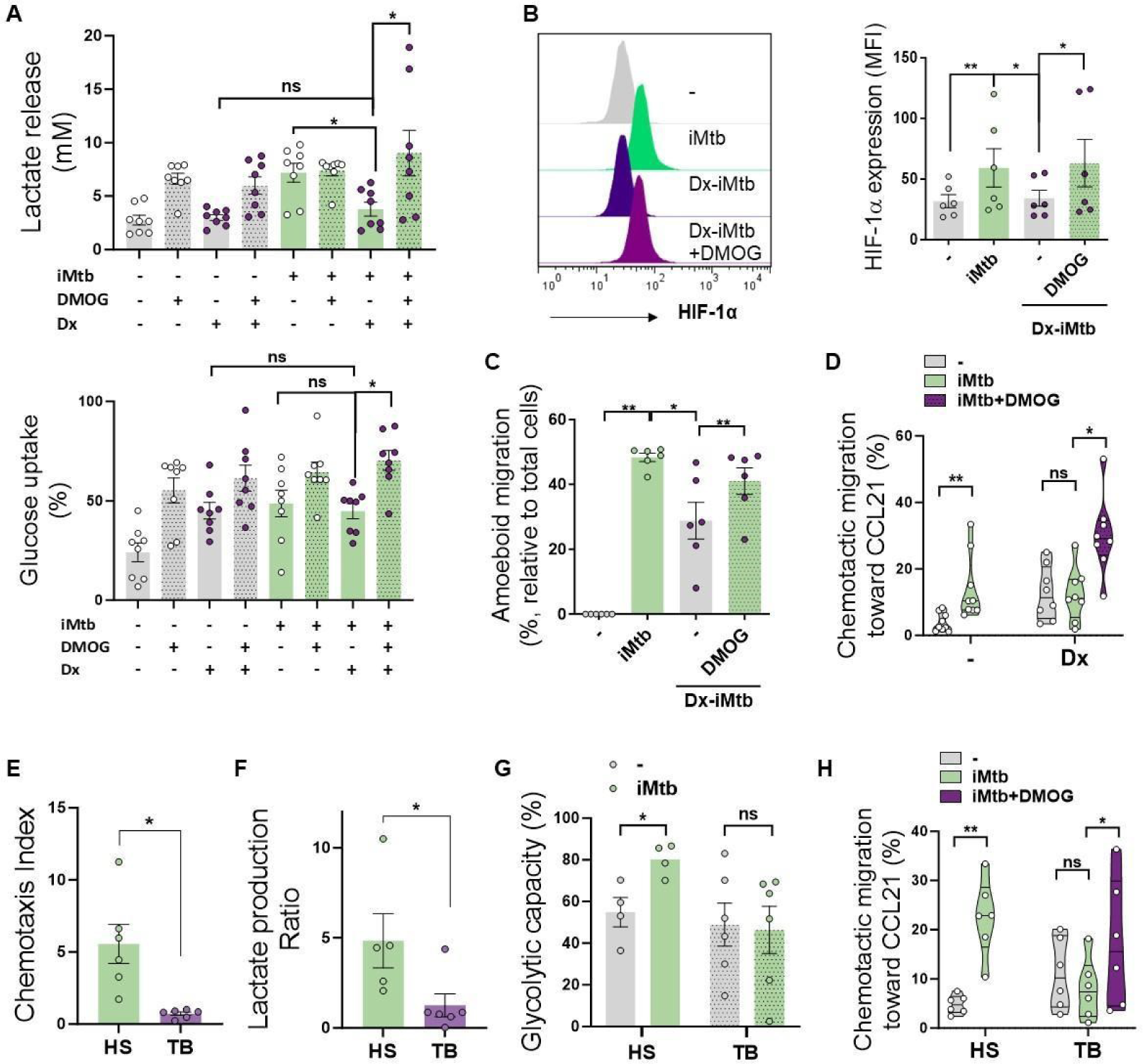
Stabilization of HIF-1α promotes migration of tolerogenic DCs and Mo-DCs from TB patients. Tolerogenic Mo-DCs were generated by dexamethasone (Dx) treatment and were stimulated (or not) with iMtb in the presence or absence of HIF-1α activator DMOG. **(A)** Lactate release and glucose uptake as measured in supernatant (N=8). **(B)** Mean fluorescence intensity (MFI) of HIF-1α. Representative histograms and quantification are shown (N=6). **(C)** Three-dimensional amoeboid migration of DCs through a collagen matrix. After 24 h of migration, images of stacks within the matrix were taken every 30 µm. Percentage of migrating cells were defined as cells in the stacks within the matrix relative to total number of cells (N=6). **(D)** Chemotactic activity towards CCL21 *in vitro* (N=6). **(E-H)** Mo-DCs were generated from healthy subjects (HS) or TB patients, and DCs were stimulated (or not) with iMtb. **(E)** Chemotaxis index towards CCL21 (relative to unstimulated DCs) (N=6). **(F)** Lactate production ratio relative to unstimulated DCs (N=6). **(G)** Glycolytic capacity assessed by SCENITH (N=4). **(H)** Chemotactic activity towards CCL21 of Mo-DCs from TB patients stimulated with iMtb and treated or not with DMOG (N=6). Statistical significance assessed by **(A-D)** 2-way ANOVA followed by Tukey’s multiple comparisons test (∗p < 0.05; ∗∗p < 0.01); **(E-G)** Unpaired T test (∗p < 0.05); **(H)** Paired T test (∗p < 0.05). The data are represented as scatter plots with each circle representing a single individual, means ± SEM are shown.

### CD16^+^ monocytes from TB patients show increased glycolytic capacity

Since we observed differences in the metabolic activity of DCs derived from monocytes of TB patients when compared to healthy donors, we next focused on evaluating the release of lactate by DC precursors from both subject groups during the first hours of DC differentiation with IL-4/GM-CSF. We found a high release of lactate by monocytes from TB patients compared to healthy donors after 1 h of differentiation (**Figure 7A**). Lactate accumulation increased in both subject groups after 24 h with IL-4/GM-CSF (**Figure 7A**). Based on these differential glycolytic activities displayed by DC precursors from both subject groups at very early stages of the differentiation process, we decided to evaluate the *ex vivo* metabolic profile of monocytes using SCENITH. To this end, we assessed the baseline glycolytic capacity of the three main populations of monocytes: classical (CD14^+^CD16^−^), intermediate (CD14^+^CD16^+^), and non-classical (CD14^dim^CD16^+^) monocytes. We found that both populations of CD16^+^ monocytes from TB patients had a higher glycolytic capacity than monocytes from healthy donors (**Figure 7B**). Moreover, the glycolytic capacity of CD16^+^ monocytes (CD14^+^CD16^+^ and CD14^dim^CD16^+^) correlates with time since the onset of TB-related symptoms (**Figure 7C**), with no association to the extent or severity of lung disease (unilateral/bilateral lesions and with/without cavities, **Figure S8**). To further expand the metabolic characterisation of monocyte subsets from TB patients, we used previously published transcriptomic data (GEO accession number: GSE185372) of CD14^+^CD16^-^, CD14^+^CD16^+^ and CD14^dim^CD16^+^ monocytes isolated from individuals with active TB, latent TB (IGRA^+^), as well as from TB negative healthy controls (IGRA^-^)^45^. Within this framework, we performed high-throughput GeneSet Enrichment Analysis (GSEA) using the BubbleMap module of BubbleGUM, which includes a multiple testing correction step to allow comparisons between the three monocyte subsets^46^. As expected, this approach reveals enrichments in genes associated with interferon responses (alpha and gamma) in patients with active TB compared to healthy donors (either IGRA^-^ or latent TB) for all three monocyte subsets (**Figure 7D**). Consistent with our findings, glycolysis increases in active TB in both CD14^+^CD16^+^ and CD14^dim^CD16^+^ monocytes (albeit not significant), while it appears to decrease in classical CD14^+^CD16^-^ monocytes (**Figure 7D**). Unlike CD14^+^CD16^-^ cells, the inflammatory response is notably enriched in CD14^+^CD16^+^ and CD14^dim^CD16^+^ monocytes from patients with active TB compared those with latent TB or healthy subjects (**Figure 7D**), suggesting that their glycolytic profile correlates with a higher inflammatory state. Finally, no significant enrichment of oxidative phosphorylation-associated genes was found in any of the performed comparisons (**Figure 7D**). Taken together, these results demonstrate that TB disease is associated with an increased activation and glycolytic profile of circulating CD16^+^ monocytes.

**Figure 7.**
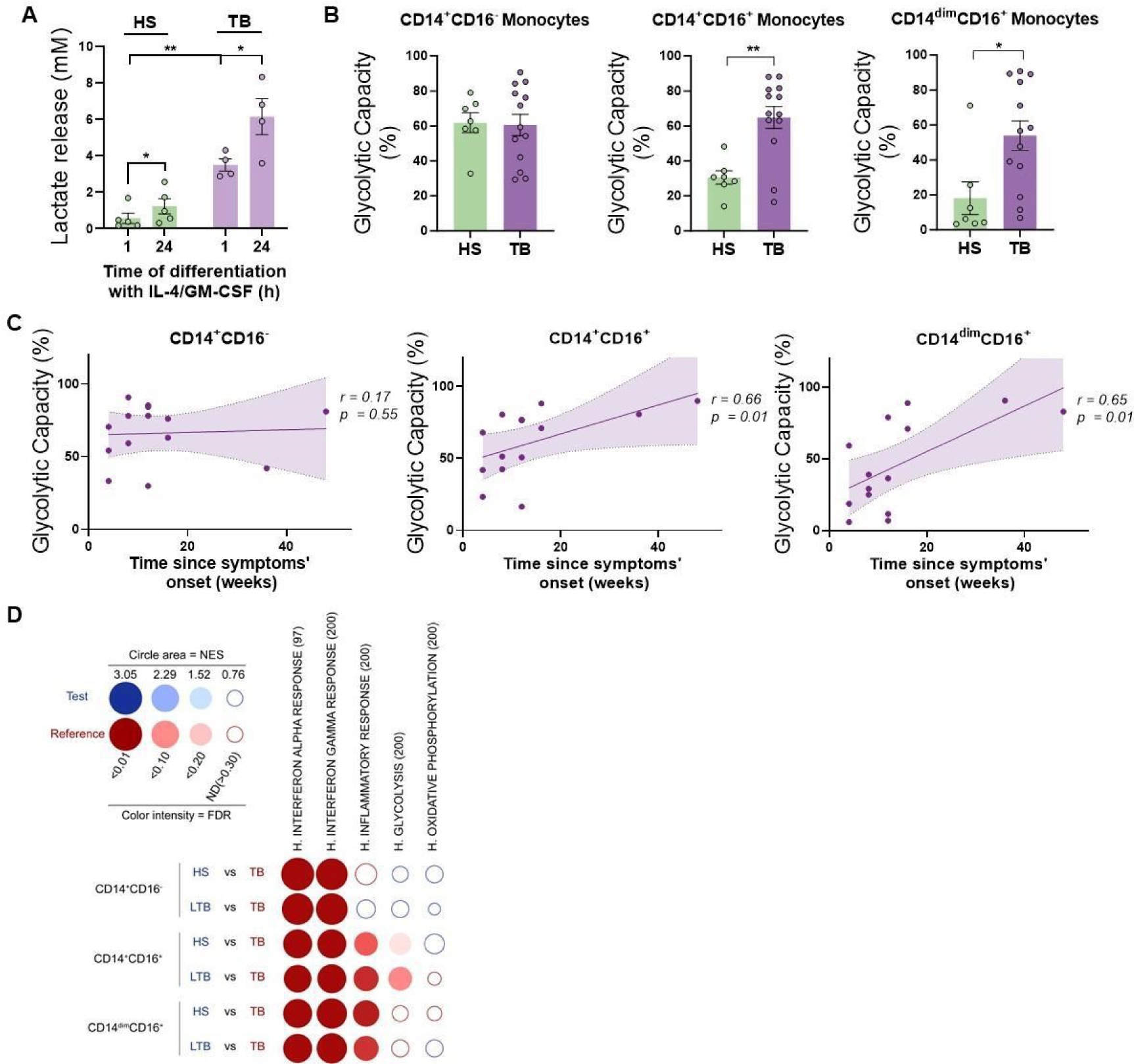
CD16^+^ monocytes from TB patients show increased glycolytic capacity. **(A)** Monocytes from TB patients or healthy subjects (HS) were isolated and cultured with IL-4 and GM-CSF for 24 h. Accumulation of lactate in culture supernatants were measured at 1 and 24 h of differentiation (N=5). (**B**) Glycolytic capacity measured by SCENITH of monocyte subsets as defined by their CD14 and CD16 expression from HS and TB patients (N=7). **(C)** Correlation analysis between the baseline glycolytic capacity and the evolution time of TB symptoms for each monocyte subset (CD14^+^CD16^-^, CD14^+^CD16^+^ and CD14^dim^CD16^+^, N=14). Linear regression lines are shown. Spearman’s rank test. The data are represented as scatter plots with each circle representing a single individual, means ± SEM are shown. **(D)** BubbleMap analysis, a high-throughput extension of GSEA, on the pairwise comparisons of monocytes from healthy patients (HS) or donors with latent TB (LTB) vs patients with active TB (TB), for each monocyte subset (CD14^+^CD16^-^, CD14^+^CD16^+^ and CD14^dim^CD16^+^). The gene sets shown come from the Hallmark (H.) collection of the Molecular Signature Database (MSigDB). The colors of the BubbleMap correspond to the population from the pairwise comparison in which the geneset is enriched (red if geneset is enriched in TB). The bubble area is proportional to the GSEA normalized enrichment score (NES). The intensity of the color corresponds to the statistical significance of the enrichment, derived by computing the multiple testing-adjusted permutation-based p-value using the Benjamini–Yekutieli correction. Enrichments with a statistical significance above 0.30 are represented by empty circles. Statistical significance was assessed by **(A)** Paired T test for 0 *vs.* 24h (∗p < 0.05) and 2-way ANOVA for HS *vs.* TB at each time (∗∗p < 0.01); **(B)** unpaired T test (∗p < 0.05; ∗∗p < 0.01).

### HIF1-α activation in CD16^+^ monocytes from TB patients leads to differentiated DCs with a poor migration capacity

Since circulating CD16^+^ monocytes from TB patients are highly glycolytic, we evaluated the expression of HIF1-α among the populations. We found that CD16^+^ monocytes from TB patients exhibited a higher expression of HIF1-α than from healthy donors (**Figure 8A**). As we have previously demonstrated that CD16^+^ monocytes from TB patients generate aberrant DCs^41^, we hypothesized that the different metabolic profile of this monocyte subset could yield DCs with some sort of exhausted glycolytic capacity and thus lower migration activity upon Mtb exposure. To test this hypothesis, we treated with DMOG to increase the activity of HIF-1α during the first 24 h of monocyte differentiation from healthy donors, leading to an exacerbated increase in lactate release at early stages of the differentiation (**Figure 8B**). Such early addition of DMOG to healthy monocytes resulted in the generation of DCs (6 days with IL-4/GM-CSF) characterized by equivalent levels of CD1a as control DCs, with a significant decrease in the expression of DC-SIGN (**Figure S9A**). In terms of activation marker expression, DCs differentiated from DMOG-pretreated cells responded to iMtb by upregulating CD86 at higher levels compared to control cells, with an accompanying trend towards reduced upregulation of CD83 (**Figure S9B**). We also observed that DCs from DMOG pretreated-cells exhibited a lower migratory capacity in response to iMtb (**Figure 8C**), reminiscent of the 2D migration capacities of Mo-DCs from TB patients. Altogether, our data suggest that the activated glycolytic status of monocytes from TB patients leads to the generation of DCs with low motility in response to Mtb.

**Figure 8.**
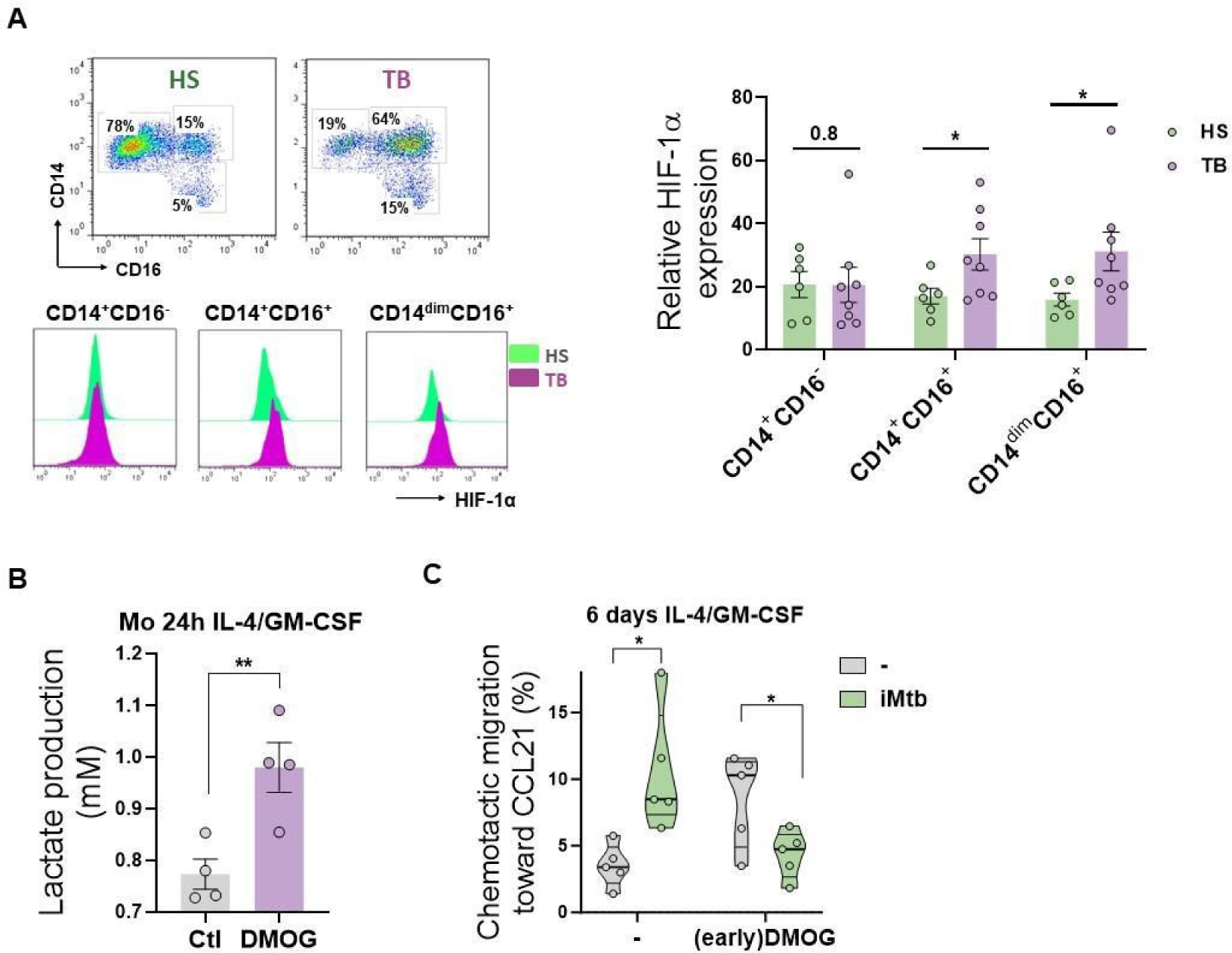
HIF1-α activation in CD16^+^ monocytes from TB patients leads to DCs with poor migration capacity. **(A)** *Ex-vivo* determination of HIF-1α expression by monocytes from healthy subjects (HS) or TB patients (TB) for each monocyte subset (CD14^+^CD16^-^, CD14^+^CD16^+^ and CD14^dim^CD16^+^) (N=6). **(B-C)** Monocytes from HS were treated with DMOG during the first 24 h of differentiation with IL-4/GM-CSF (earlyDMOG) and removed afterwards. On day 6 of differentiation, cells were stimulated (or not) with iMtb. **(B)** Monocyte lactate release after 24 h of DMOG addition (N=4). **(C)** Chemotactic activity towards CCL21 of DCs (N=5). Statistical significance was assessed by **(B)** paired T test (∗∗p < 0.01); **(C)** 2-way ANOVA followed by Tukey’s multiple comparisons test (∗p < 0.05). The data are represented as scatter plots with each circle representing a single individual, means ± SEM are shown.

## Discussion

In this study, we provide evidence for the role of HIF-1α-mediated glycolysis in promoting the migratory capacity of DCs upon encounter with iMtb. Our approach to quantify the *ex vivo* metabolism of monocytes shows that CD16^+^ monocytes from TB patients display an exacerbated glycolytic activity that may result in the generation of DCs with poor migratory capacities in response to iMtb. Our results suggest that under extensive chronic inflammatory conditions, such as those found in TB patients, circulating monocytes may be metabolically preconditioned to differentiate into DCs with low migratory potential.

Upon Mtb infection of naive mice, initial accumulation of activated CD4^+^ T cells in the lung is delayed, occurring between 2-3 weeks post-infection^3,47^. The absence of sterilizing immunity induced by TB vaccines, such as BCG, has been proposed to result from delayed activation of DCs and the resulting delay in antigen presentation and activation of vaccine-induced CD4^+^ T-cell responses^48^. In this context, it was demonstrated that Mtb-infected Mo-DCs recruited to the site of infection exhibit low CCR7 expression and impaired migration to lymph nodes compared to uninfected Mo-DCs^49^. Additionally, Mo-DCs have been found to play a key role in transporting Mtb antigens from the lung to the draining lymph node, where conventional DCs present antigens to naive T cells^50^. The migratory capacity of responding DCs is thus of paramount importance to the host response to Mtb infection.

Here, we found that Mtb exposure triggers glycolysis in Mo-DCs from healthy donors, which promotes their migration capacity in a HIF-1α-dependent manner. Recently, it was shown that glycolysis was required for CCR7-triggered murine DC migration in response to LPS^22–24^. Glycolysis was also reported to be required for the migration of other immune cells such as macrophages^51^ and regulatory T cells^52^. Consistently, we show that inhibition of HIF-1α-dependent glycolysis impairs human Mo-DC migration upon Mtb stimulation. The link between cellular metabolism and migratory behavior are supported by studies that have elucidated how glycolysis can be mechanically regulated by changes in the architecture of the cytoskeleton, ultimately impacting the activity of glycolytic enzymes^53,54^. In addition, interesting links between cellular mechanics and metabolism have been previously described for DCs, highlighting the potential to alter DC mechanics to control DC trafficking and consequently T cell priming^16^. However, studies focused on the molecular mechanisms by which metabolic pathways impact the machinery responsible for cell movement in the context of TB infection will be required to better understand and design therapeutic manipulation.

Our research indicates that DCs exhibit upregulated glycolysis following stimulation or infection by Mtb. This metabolic shift is crucial for facilitating cell migration to the draining lymph nodes, an essential step in mounting an effective immune response. Yet, it remains uncertain whether this glycolytic induction reaches a threshold conducive to generating a protective immune response, a matter that our findings do not definitively address. In addition, we demonstrated that tolerogenic DCs induced by dexamethasone as well as DCs derived from TB patient monocytes exhibit lower lactate release and impaired trafficking toward CCL21 upon Mtb stimulation; both phenotypes could be rescued by stabilization of HIF-1α expression. To our knowledge, this is the first study to address how the metabolic status of monocytes from TB patients influences the migratory activity of further differentiated DCs. According to our findings, the activation status of the glycolysis/HIF-1α axis in monocytes would be a predictor of refractoriness to differentiation into migratory DCs in TB. With respect to the metabolism of tolerogenic DCs broadly, our results are consistent with reported data showing that DC tolerance can be induced by drugs promoting OXPHOS, such as vitamin D and dexamethasone^55–57^. It was interesting to note that, although migration of tolerogenic DCs did not increase upon Mtb stimulation, it was increased under basal conditions, which agrees with previous data showing a high steady-state migration capacity of putatively tolerogenic DCs^58^.

It has been widely demonstrated that immune cells can switch to glycolysis following engagement of TLRs^59^. Our work showed that TLR2 ligation by either viable or irradiated Mtb was necessary to trigger glycolysis in DCs, at least at early times post-stimulation. In fact, even bystander DCs increased their glycolytic activity in Mtb-infected cultures, suggesting that mycobacterial antigens or bacterial debris present in the microenvironment may be sufficient to trigger TLR-dependent glycolysis. In the context of natural infection *in vivo*, we foresee that DC with different levels of infection will coexist, some with low bacillary load that, according to our data, may be able to trigger glycolysis and migrate, while others highly infected DCs would more likely die^60^. It remains to be elucidated whether persistent interaction between DCs and Mtb might lead to an attenuation in glycolysis over time, as has been reported for macrophages^61^. In this regard, our data demonstrates that chronic Mtb infection leads to monocytes bearing an exacerbated glycolytic status likely tied to prolonged and or excessive stimulation of membrane bound TLRs in circulation, which results in DCs with an exhausted glycolytic capacity. Although DCs stimulated with iMtb in the presence of a HIF-1α inhibitor exhibited differences in activation markers and cytokine profile, we found that they were still able to activate CD4^+^ T cells from PPD+ donors in response to iMtb. These findings complement previous evidence showing that LPS-induced mature DCs inhibit T-cell responses through HIF-1α activation in the presence of glucose, leading to greater T cell activation capacity in low glucose contexts such as at the interface between DCs and T cells^25^. In this work, we did not detect an impact on T cell activation upon HIF-1α inhibition in DCs, but we observed a clear reduction in their migration capacity that may limit or delay DC encounters with T cells *in vivo*, leading to poor T cell activation in the lymph nodes. In this regard, mouse studies have shown that DC migration directly correlates with T cell proliferation^62^. However, we cannot rule out the possibility that other CD4^+^ T cell subsets (such as regulatory T cells), CD1-restricted T cells, and/or CD8^+^ T cell subsets could be differentially activated by iMtb-stimulated DCs lacking HIF-1α activity.

Three different populations of human monocytes have been identified: classical (CD14^+^, CD16^−^), intermediate (CD14^+^, CD16^+^), and non-classical (CD14^dim^, CD16^+^) monocytes^63^. These monocyte subsets are phenotypically and functionally distinct. Classical monocytes readily extravasate into tissues in response to inflammation, where they can differentiate into macrophage-like or DC-like cells^64^; intermediate monocytes are well-suited for antigen presentation, cytokine secretion, and differentiation; and non-classical monocytes are involved in complement and Fc gamma-mediated phagocytosis and their main function is cell adhesion^65,66^. Unlike non-classical monocytes, the two CD14^+^ monocyte populations are known to extravasate into tissues and thus are likely to act as precursors capable of giving rise to Mo-DCs in inflamed tissues. However, the DC differentiation capacity of the intermediate population is still not well defined. We previously demonstrated that monocytes from TB patients generate aberrant DCs, and that CD16^+^ monocytes generate aberrant DCs upon treatment with GM-CSF and IL-4^41^. Here, we demonstrated that glycolysis seems to play a dual role during DC differentiation from monocytes, on the one side, being required for fully differentiated-DC migration to lymph nodes in response to Mtb and, on the other side, leading to DCs with poor iMtb-responsive migratory capacity if activated during the onset of DC differentiation. In this regard, DCs from healthy subjects respond to iMtb by inducing a glycolytic and migratory profile, while monocytes isolated from TB patients exhibit an unusual early glycolytic state that results in the ulterior generation of DCs with low glycolytic and migratory activities in response to Mtb. Similarly, we found that CD16^+^ cells from TB patients display an activated glycolytic status, as well as elevated HIF-1α expression levels compared to their healthy counterparts. Additionally, we showed that monocytes from TB patients are not only enriched in CD16^+^ cells, but also display an altered chemokine receptor expression profile^67^, demonstrating that both phenotype and function of a given monocyte subset may differ under pathological conditions. While it is difficult to determine whether the heightened glycolytic profile of monocytes may limit their differentiation into DCs *in vivo*, we provided evidence that an increase in HIF1α-mediated glycolysis in precursors leads to the generation of cells with poor ability to migrate in response to CCL21 *in vitro.* In line with this observation, a recent study revealed a significant increase in the glycolytic capacity occurs during the first 24 h of monocyte differentiation towards a tolerogenic DC phenotype, as induced by vitamin D3^26^, highlighting the detrimental role of an early activated inflammatory profile in DC precursors. A possible explanation for these effects may be found in lactate accumulation in monocytes during DC differentiation. Lactate signaling in immune cells leads to metabolic alterations in DCs that program them to a regulatory state^68^, and lactate has also been shown to suppress DC differentiation and maturation^18^; thus, excessive precursor glycolytic activity may result in DCs biased toward regulatory functions.

Taken together, our data offer new insights into the immunometabolic pathways involved in the trafficking of DCs to the lymph nodes. These insights may have various implications depending on factors such as timing, cell type, and location induction of the HIF1α/glycolysis axis. On the one hand, nurturing HIF-1α-mediated glycolytic activity in DCs during the early stages of infection could potentially enhance the effectiveness of preventive strategies for TB. Particularly noteworthy is the significant impact revealed in studies where the number of DCs reaching the lymph node proved to be a crucial factor in determining the success of DC-based vaccination^62^. On the other hand, premature activation of glycolysis in precursors, as observed in CD16+ monocytes from severe TB patients, could disrupt the delicate balance necessary for an optimal immune response. This variability is consistent with the paradigm of “too much, too little,” as demonstrated by the dual roles of IFNγ^69^ and TNFα^70^ in the context of TB. It also underscores the vital importance of maintaining an equilibrium in inflammatory responses. This study lays the foundation for further exploration into the potential systemic impact of the HIF1α/glycolysis axis within the realm of chronic inflammation contrasting with its role in a local setting during the acute phase of infection. By enhancing our understanding, these findings aim to guide the development of innovative preventive and therapeutic strategies for TB.

## Methods

### Chemical Reagents

LPS from Escherichia coli O111:B4 was obtained from Sigma-Aldrich (St. Louis, MO, USA). Dexamethasone (Dx) was from Sidus (Buenos Aires, Argentina). PX-478 2HCL was purchased from Selleck Chemicals (Houston, USA) and DMOG from Santa Cruz, Biotechnology (Palo Alto, CA, USA). Additionally, GSK2837808A was purchased from Cayman Chemical (Michigan, USA) together with echinomycin and sodium oxamate.

### Bacterial strain and antigens

Mtb H37Rv strain was grown at 37°C in Middlebrook 7H9 medium supplemented with 10 % albumin-dextrose-catalase (both from Becton Dickinson, New Jersey, USA) and 0.05 % Tween-80 (Sigma-Aldrich). The Mtb γ-irradiated H37Rv strain (NR-49098) was obtained from BEI Resource (NIAID, NIH, USA). The RFP-expressing Mtb strain was gently provided by Dr. Fabiana Bigi (INTA, Castelar, Argentina).

### Preparation of monocyte-derived DCs

Buffy coats from healthy donors were prepared at Centro Regional de Hemoterapia Garrahan (Buenos Aires, Argentina) according to institutional guidelines (resolution number CEIANM-664/07). Informed consent was obtained from each donor before blood collection. Monocytes were purified by centrifugation on a discontinuous Percoll gradient (Amersham, Little Chalfont, UK) as previously described^71^. Then, monocytes were allowed to adhere to 24-well plates at 5×10^5^ cells/well for 1 h at 37°C in warm RPMI-1640 medium (ThermoFisher Scientific, Waltham, MA). The mean purity of adherent monocytes was 85 % (range: 80-92 %). The medium was then supplemented to a final concentration of 10 % Fetal Bovine Serum (FBS, Sigma-Aldrich), human recombinant Granulocyte-Macrophage Colony-Stimulating Factor (10 ng/ml, GM-CSF, Peprotech, New Jersey, USA) and IL-4 (20 ng/ml, Biolegend, San Diego, USA). Cells were allowed to differentiate for 5-7 days (DC-SIGN^+^ cells in the culture > 90 %).

### DC stimulation

DCs were stimulated with either irradiated Mtb (iMtb) or viable Mtb at equivalent OD_600_ doses for 24 h at 37 °C. The cells were washed three times, and their phenotype and functionality were evaluated together with survival of activated cells; cell number and viability were determined by either trypan blue exclusion assays or MTT. Infections were performed in the biosafety level 3 (BSL-3) laboratory at the Unidad Operativa Centro de Contención Biológica (UOCCB), ANLIS-MALBRAN (Buenos Aires), according to the biosafety institutional guidelines.

### DC treatments

When indicated, neutralizing monoclonal antibodies (mAb), or their corresponding isotype antibodies as mock controls, were added 30 min prior to DC stimulation to inhibit TLR2 (309717, Biolegend) or TLR4 (312813, Biolegend). In addition, DCs were incubated with PX-478 (20 µM) or Echinomycin (1 nM) with the purpose of inhibiting HIF-1α activity, DMOG (50 µM) to stabilize HIF-1α, and oxamate (20 mM) or GSK2837808A (20 µM) to inhibit LDH. DC stimulation with iMtb occurred 30 min after treatment without drug washout.

In figure 6 and S6, dexamethasone-induced tolerogenic dendritic cells (Dx-DC) were generated by incubating DCs with 0.1 µM of dexamethasone for 1 h. Thereafter, cells were washed, and “complete medium” was added. Tolerogenic Dx-DCs were then stimulated (or not) with iMtb in the presence or not of DMOG (50 µM).

### Determination of metabolite concentrations

Lactate production and glucose concentrations in the culture medium was measured using the spectrophotometric assays Lactate Kit and Glicemia Enzimática AA Kit both from Wiener (Argentina), which are based on the oxidation of lactate or glucose, respectively, and the subsequent production of hydrogen peroxide^72^. The consumption of glucose was determined by assessing the reduction in glucose levels in culture supernatants in comparison with RPMI 10 % FBS. The absorbance was read using a Biochrom Asys UVM 340 Microplate Reader microplate reader and software.

### Quantitative RT-PCR

Total RNA was extracted with Trizol reagent (Thermo Fisher Scientific) and cDNA was reverse transcribed using the Moloney murine leukemia virus reverse transcriptase and random hexamer oligonucleotides for priming (Life Technologies, CA, USA). The expression of the genes *HIF-1α* and *LDH-A* was determined using the PCR SYBR Green sequence detection system (Eurogentec, Seraing, Belgium) and the CFX Connect Real-Time PCR Detection System (Bio-Rad, CA, USA). Gene transcript numbers were standardized and adjusted relative to eukaryotic translation elongation factor 1 alpha 1 (*EeF1A1*) transcripts. Gene expression was quantified using the ΔΔCt method. Primers used for RT-PCR were as follows: EeF1A1 Fwd: 5’-TCGGGCAAGTCCACCACTAC −3’ and Rev: 5’-CCAAGACCCAGGCATACTTGA-3’; HIF-1α Fwd: 5’-ACTAGCCGAGGAAGAACTATGAA-3’ and Rev: 5’-TACCCACACTGAGGTTGGTTA-3’; and LDH-A Fwd: 5’-TGGGAGTTCACCCATTAAGC-3’ and Rev: 5’-AGCACTCTCAACCACCTGCT-3’.

### Immunofluorescence analysis

FITC-, PE- or PerCP.Cy5.5-labelled mAbs were used for phenotypic analysis of the following cell-surface receptor repertoires: FITC-anti-CD1a (clone HI149, eBioscience), PE-anti-DC-SIGN (clone 120507, RD System), PerCP.Cy5.5-anti-CD86 (clone 374216, Biolegend), FITC-anti-CD83 (clone HB15e, eBioscience), PE-anti-PD-L1 (clone MIH1, BD Pharmingen) and in parallel, with the corresponding isotype control antibody. Approximately 5×10^5^ cells were seeded into tubes and washed once with PBS. Cells were stained for 30 min at 4°C and washed twice. Additionally, cells were stained for 40 min at 4°C with fluorophore-conjugated antibodies PE-anti-Glut1 (clone 202915 R&D Systems, Minnesota, USA) and in parallel, with the corresponding isotype control antibody. For HIF-1α determination, DCs were permeabilized with methanol and incubated with PE-anti-HIF-1α (clone 546-16, Biolegend). Stained populations were gated according to forward scatter (FSC) and side scatter (SSC) analyzed on FACScan (Becton Dickinson). Isotype matched controls were used to determine auto-fluorescence and non-specific staining. Analysis was performed using the FCS Express (De Novo Software) and results were expressed as median fluorescence intensity (MFI) or percentage of positive cells.

### Soluble cytokines determinations

Supernatants from DC populations or DC-T cell cocultures were harvested and assessment of TNF-α, IL-10, IL-17A or IFN-γ production was measured by ELISA, according to manufacturers’ instructions (eBioscience). The detection limit was 3 pg/ml for TNF-α and IL-17A, 6 pg/ml for IFN-γ, and 8 pg/ml for IL-10.

### CD4^+^ T cell activation assay

Specific lymphocyte activation (recall) assays were carried out in cells from tuberculin purified protein derivative-positive skin test (PPD^+^) healthy donors by culturing DC populations and autologous T cells at a ratio of 10 T cells to 1 DC in round bottom 96-well culture plates for 5 days as detailed previously^9^. The numbers of DCs were adjusted to live cells before the start of the co-cultures. After 5 days, CD4^+^ T cell subsets were identified by immunolabeling according to the differential expression of CCR4, CXCR3, and CCR6 as previously reported^73^: CXCR3^+^CCR4^−^CCR6^−^ (Th1), CXCR3^−^CCR4^+^CCR6^−^ (Th2), CXCR3^−^CCR4^+^CCR6^+^ (Th17), and CXCR3^+^CCR4^−^CCR6^+^ (Th1* or Th1/Th17). The fluorochrome-conjugated antibodies used for flow cytometry analysis were CD4-FITC (clone A161A1, Biolegend), CXCR3-PE-Cy7 (clone G025H7, Biolegend), CCR4-PerCPCy5.5 (clone 1G1, BD Bioscience), CCR6-APC (clone 11A9, BD Bioscience), and CD3-APC-Cy7 (clone HIT3a, Biolegend). A viability dye, Zombie Violet (Biolegend), was used to exclude dead cells. Fluorescence Minus One (FMO) control was used to set proper gating for CXCR3-PE-Cy7, CCR4-PerCPCy5.5, and CCR6-APC detection. Cells were analyzed by fluorescence-activated cell sorting (FACS), using the BD FACSCANTO cytometer and FlowJo^TM^ Software (BD Life Sciences).

### Chemotactic activity of DCs

Each DC population (4×10^5^ cells in 75 µl) was placed on the upper chamber of a transwell insert (5 µm pore size, 96-well plate; Corning), and 230 µL of media (RPMI with 0.5 % FCS) with human recombinant CCL21 (200 ng/ml) (Peprotech) was placed in the lower chamber. After 3 h, cells that had migrated to the lower chamber were removed and analyzed. The relative number of cells migrating was determined on a flow cytometer using Calibrite beads (BD Biosciences), where a fixed number of beads was included in each sample and the number of cells per 1,000 beads was evaluated. Data were normalized to the number of initial cells.

### *In vivo* migration assay

DCs were differentiated from bone marrow precursors obtained from naive BALB/c mice in the presence of murine GM-CSF (10 ng/ml) and IL-4 (10 ng/ml) both from Biolegend for 7 days. After differentiation, DCs were treated with oxamate (20 mM) or PX-478 (10 uM) and stimulated with iMtb. After 24 h, DCs were stained with CFSE (5 µM) and inoculated intradermally in the inguinal zone of naïve BALB/c mice. 3 h post-injection, inguinal lymph nodes close to the site of inoculation were harvested and cells were stained with fluorophore-conjugated antibody PE-anti-CD11c (clone HL3, BD Pharmingen). Analysis was performed using the FlowJo Software and results were expressed as the percentage of CFSE^+^/CD11c^+^ cells.

### 3D migration assay

0.5×10^5^ DCs were seeded on top of fibrillar collagen matrices polymerized from Nutragen 2 mg/ml, 10 % v/v MEM 10X (MEM invitrogen, Carlsbad, CA), UltraPure distilled water and 4-6 % v/v bicarbonate buffer (pH=9) 7.5 %. After 24 h, cellular migration was quantified by taking images using an inverted microscope (Leica DMIRB, Leica Microsystems, Deerfield, IL) and the software Metamorph, as described previously^74^. Alternatively, in a similar manner, matrices were polymerized using Collagen (Sigma-Aldrich, C9791-10MG) in figure 5C and D. After 24 h of cellular migration, matrices were fixed with paraformaldehyde (PFA) 4 % during 30 min at room temperature and stained with DAPI (Cell signaling). Collagen was removed and membranes were mounted with DAKO. Images were taken by confocal microscopy (FluoView FV 1000), and cells were counted per field.

### Measurement of cell respiration with Seahorse flux analyzer

Bioenergetics were determined using a seahorse XFe24 analyzer. ATP production rates and relative contribution from the glycolysis and the OXPHOS were measured by the Seahorse XF Real-Time ATP Rate Assay kit. DCs (2×10^5^ cells/well) were cultured in 3 wells per condition. The assay was performed in XF Assay Modified DMEM. Three consecutive measurements were performed under basal conditions and after the sequential addition of oligomycin and rotenone/antimycin (Agilent, USA). Extracellular acidification rate (ECAR) and oxygen consumption rate (OCR) were measured. Mitochondrial ATP production rate was determined by the decrease in the OCR after oligomycin addition. On the other hand, the complete inhibition of mitochondrial respiration with rotenone plus antimycin A, allows accounting for mitochondrial-associated acidification, and when combined with proton efflux rate (PER) data, allows calculation of Glycolysis ATP production rate. All OCR and ECAR values were normalized. Briefly, before the assay brightfield imaging was performed. Cellular area per condition was calculated by ImageJ software and imported into Wave (Agilent) using the normalization function.

### SCENITH™ assay

SCENITH experiments were performed as previously described^32^ using the SCENITH kit containing all reagents and anti-puromycin antibodies (www.scenith.com). Briefly, DCs or PBMCs were treated for 40 min at 37°C in the presence of the indicated inhibitors of various metabolic pathways and puromycin. After the incubation, puromycin was stained using a fluorescently-labeled anti-Puromycin monoclonal antibody (clone R4743L-E8) with Alexa Fluor 647 or Alexa Fluor 488, and analyzed by flow cytometry. For metabolic analysis of monocyte subsets, PBMCs were labeled with PE-anti-CD16 (clone 3G8, Biolegend) and PECy7-anti-CD14 (clone HCD14, Biolegend) mAbs. The impact of the various metabolic inhibitors was quantitated as described^32^.

### Transmission electron microscopy

DCs were fixed in 2.5 % glutaraldehyde / 2 % PFA (EMS, Delta-Microscopies) dissolved in 0.1 M Sorensen buffer (pH 7.2) for 1 h at room temperature, and then preserved in 1 % PFA dissolved in Sorensen buffer. Adherent cells were treated for 1 h with 1 % aqueous uranyl acetate then dehydrated in a graded ethanol series and embedded in Epon. Sections were cut on a Leica Ultracut microtome and ultrathin sections were mounted on 200 mesh onto Formvar carbon coated copper grids. Finally, thin sections were stained with 1 % uranyl acetate and lead citrate and examined with a transmission electron microscope (Jeol JEM-1400) at 80 kV. Images were acquired using a digital camera (Gatan Orius). For mitochondrial morphometric analysis, TEM images were quantified with the ImageJ “analyze particles” plugin in thresholded images, with size (μm^2^) settings from 0.001 to infinite. For quantification, 8–10 cells of random fields (1000x magnification) per condition were analyzed.

### Changes of mitochondrial mass

Mitochondrial mass was determined in DCs by fixing the cells with PFA 4% and labeling them with the probe MitoSpy Green FM (Biolegend). Green fluorescence was analyzed by flow cytometry (FACScan, BD Biosciences).

### GeneSet Enrichment Analysis of human monocytes

BubbleMap analysis was performed with 1,000 geneset-based permutations, and with “Signal2Noise” as a metric for ranking the genes. The results are displayed as a BubbleMap, where each bubble is a GeneSet Enrichment Analysis (GSEA) result and summarizes the information from the corresponding enrichment plot. The color of the Bubble corresponds to the population from the pairwise comparison in which the geneset is enriched. The bubble area is proportional to the GSEA normalized enrichment score (NES). The intensity of the color corresponds to the statistical significance of the enrichment, derived by computing the multiple testing-adjusted permutation-based p-value using the Benjamini–Yekutieli correction. Enrichments with a statistical significance above 0.30 are represented by empty circles.

### Patient blood donors

TB patients were diagnosed at the División Tisioneumonología, Hospital F.J.Muñiz (Buenos Aires, Argentina) by the presence of recent clinical respiratory symptoms, abnormal chest radiography, and positive culture of sputum or positive sputum smear test for acid-fast-bacilli. Written, informed consent was obtained according to the Ethics Committee from the Hospital Institutional Ethics Review Committee. Exclusion criteria included HIV positive patients and the presence of concurrent infectious diseases or comorbidities. Blood samples were collected during the first 15 days after commencement of treatment. All tuberculous patients had pulmonary TB (Table I). The term symptoms evolution refers to the time period during which a patient experiences cough and phlegm for more than 2-3 weeks, with (or without) sputum that may (or not) be bloody, accompanied by symptoms of constitutional illness (*e.g.,* loss of appetite, weight loss, night sweats, and general malaise).

**Table I.** Demographic and clinical characteristics of TB patients.

|  |  |
| --- | --- |
| **Age, years (range)** | 36 (19-67) |
| **Gender, % (number/total)** | M, 81% (31/38)<br>F, 19% (7/38) |
| **Nationality, % (number/total)** | Argentina, 76.31% (29/38)<br>Bolivia, 15.78% (6/38)<br>Paraguay, 2.63% (1/38)<br>Peru, 5.26% (2/38) |
| **TB disease localization, % (number/total)** | Pulmonary, 94% (36/38)<br>Pulmonary + Extrapulmonary, 6% (2/38) |
| **AFB\* in sputum, % (number/total)** | 3+, 21% (8/38)<br>2+, 13% (5/38)<br>1+, 52% (20/38)<br>-, 13% (5/38) |
| **Leukocyte count, mean ± SEM, cell/μL** | 8483 ± 509 |
| **Lymphocyte mean ± SEM, %** | 19 ± 2 |
| **Monocyte mean ± SEM, %** | 7 ± 0,5 |

\* Acid-fast-bacilli (AFB) in sputum: -, 1+, 2+, 3+ are defined according to the International Union Against Tuberculosis and Lung Disease (IUATLD)/ World Health Organization (WHO) quantification scale

### Statistics

All values are presented as the median ± SEM of 3-13 independent experiments. Each independent experiment corresponds to 1 donor. For Seahorse assays, OCR and PER values are shown as mean ± SD. Comparisons between unpaired experimental conditions were made using either ANOVA for parametric data or Friedman test for non-parametric data followed by Dunn’s Multiple Comparison Test. Comparisons between paired experimental conditions were made using the two-tailed Wilcoxon Signed Rank Test for non-parametric data or T test for parametric data. Correlation analyses were determined using the Spearman’s rank test. A p-value of 0.05 was considered significant.

### Study approval

#### Human specimens

The study design was reviewed and approved by the Ethics Committees of the Academia Nacional de Medicina (49/20/CEIANM) and the Muñiz Hospital, Buenos Aires, Argentina (NI #1346/21). All participants voluntarily enrolled in the study by signing an informed consent form after receiving detailed information about the research study.

#### Mouse studies

All experimental protocols were approved by the Institutional Animal Care and Use of the Experimentation Animals Committee (CICUAL number 090/2021) of the Institute of Experimental Medicine (IMEX, Buenos Aires).

## AUTHOR CONTRIBUTIONS

Conceptualization & methodology: MM, FF, MFQ, MV, TPVM, CV and LB. Investigation: MM, MJ, ZV, JB, JLMF, MG, SM, MBV, FF, VGP, MV, CV and LB. Resources: SI, RM, LC, PGM, DP, RA, ON, CV, GL-V, MdCS and LB. Formal Analysis: MM, TPVM, and LB. Writing: MM, GL-V, RA, MdCS, ON, CV, TPVM and LB. Visualization: MM, FF, CV, TPVM and LB. Funding acquisition: ON, GL-V, RA, CV, and LB. Corresponding author: LB is responsible for ownership and responsibility that are inherent to aspects of this study.

## ACKNOWLEDGMENTS

We thank the staff of the Regional Center of Hemotherapy of the Garrahan Hospital (Buenos Aires). We greatly thank Claire Lastrucci for designing the graphical abstract. In addition, we are grateful for the editing service provided by Life Science editors.

This work was supported by the Argentinean National Agency of Promotion of Science and Technology (PICT-2019-01044 and PICT-2020-00501 to LB); the Argentinean National Council of Scientific and Technical Investigations (CONICET, PIP 11220200100299CO to LB); the Centre National de la Recherche Scientifique, *Université Paul Sabatier*, the *Agence Nationale de Recherche sur le Sida et les hépatites virales (ANRS)* (ANRS2018-02, ECTZ 118551/118554, ECTZ 205320/305352, ANRS ECTZ103104 and ECTZ101971 to CV, ON and GL-V); and the French ANR JCJC-Epic-SCENITH ANR-20-CE14-0028 and CoPoC Inserm-transfert MAT-PI-17493-A-04 to RA. The funders had no role in study design, data collection, and analysis, decision to publish, or preparation of the manuscript.

