## Supplementary figures1-9 for "Elevated glycolytic metabolism of monocytes limits the generation of HIF-1α-driven migratory dendritic cells in tuberculosis"

Mariano Maio *et al.*

**This PDF file includes:**

Figs. S1 to Main Fig. 2
Figs. S2 to Main Fig. 3
Figs. S3 to Main Fig. 4
Figs. S4 to Main Fig. 4
Figs. S5 to Main Fig. 4
Figs. S6 to Main Fig. 5
Figs. S7 to Main Fig. 6
Figs. S8 to Main Fig. 7
Figs. S9 to Main Fig. 8

**Fig. S1.**

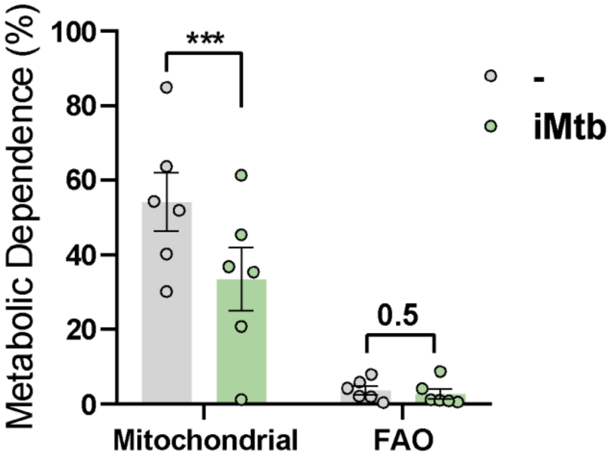


**Contribution of the fatty acid oxidation (FAO) to DC metabolism in response to irradiated Mtb**. Relative contributions of mitochondrial and FAO dependences to overall DC metabolism analyzed with SCENITH in DCs exposed or not to iMtb (N=6). Paired t test (∗∗∗$p < 0.001$) as depicted by lines. The data are represented as scatter plots with each circle representing a single individual, means ± SEM are shown.

**Fig. S2.**

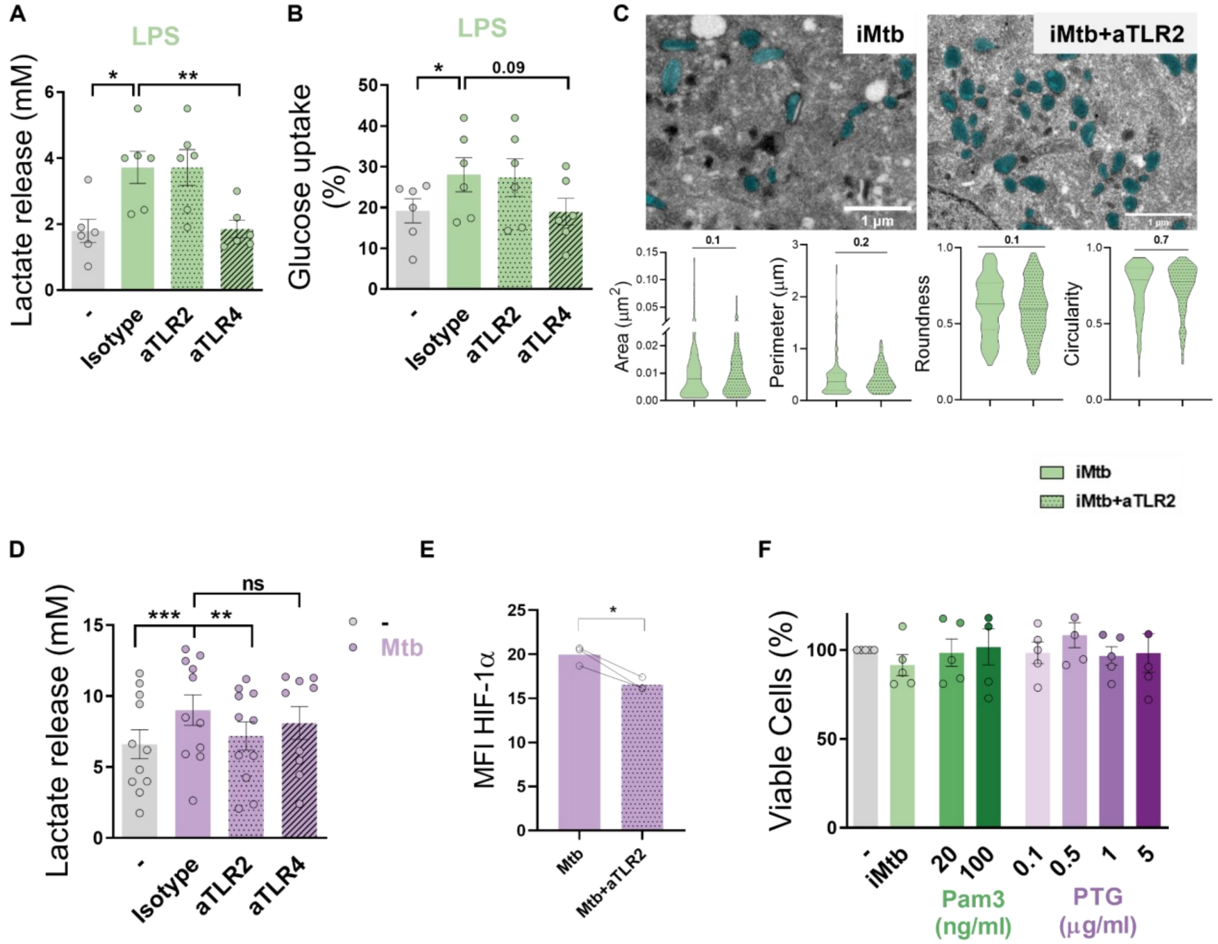


**TLR2 ligation triggers glycolysis in Mo-DCs. (A-B)** Mo-DCs were stimulated or not with LPS in the presence of neutralizing antibodies against either TLR2 (aTLR2), or TLR4 (aTLR4). **(A)** Lactate release measured in supernatant (N=6). **(B)** Glucose uptake measured in supernatant (N=6). **(C)** Morphometric analysis of mitochondria of Mo-DCs stimulated or not with iMtb in the presence of neutralizing antibodies against TLR2 (N=4). **(D-E)** Mo-DCs were infected or not with Mtb in the presence of neutralizing antibodies against either TLR2 (aTLR2), or TLR4 (aTLR4). (D) Lactate release measured in supernatant (N=11). **(E)** Mean fluorescence intensity (MFI) of HIF-1α as measured by flow cytometry (N=3). **(F)** Percentage of viable cells stimulated with the synthetic TLR2 ligand (Pam3), or different doses of the mycobacterial peptidoglycan (PTG) relative to untreated DCs (N=5). **(A-B, D-F)** 2-way ANOVA followed by Tukey's multiple comparisons test ($*p < 0.05$; $**p < 0.01$), as depicted by lines. **(C, E)** Paired t test ($*p < 0.05$). The data are represented as scatter plots with each circle representing a single individual, means ± SEM are shown.

**Fig. S3.**

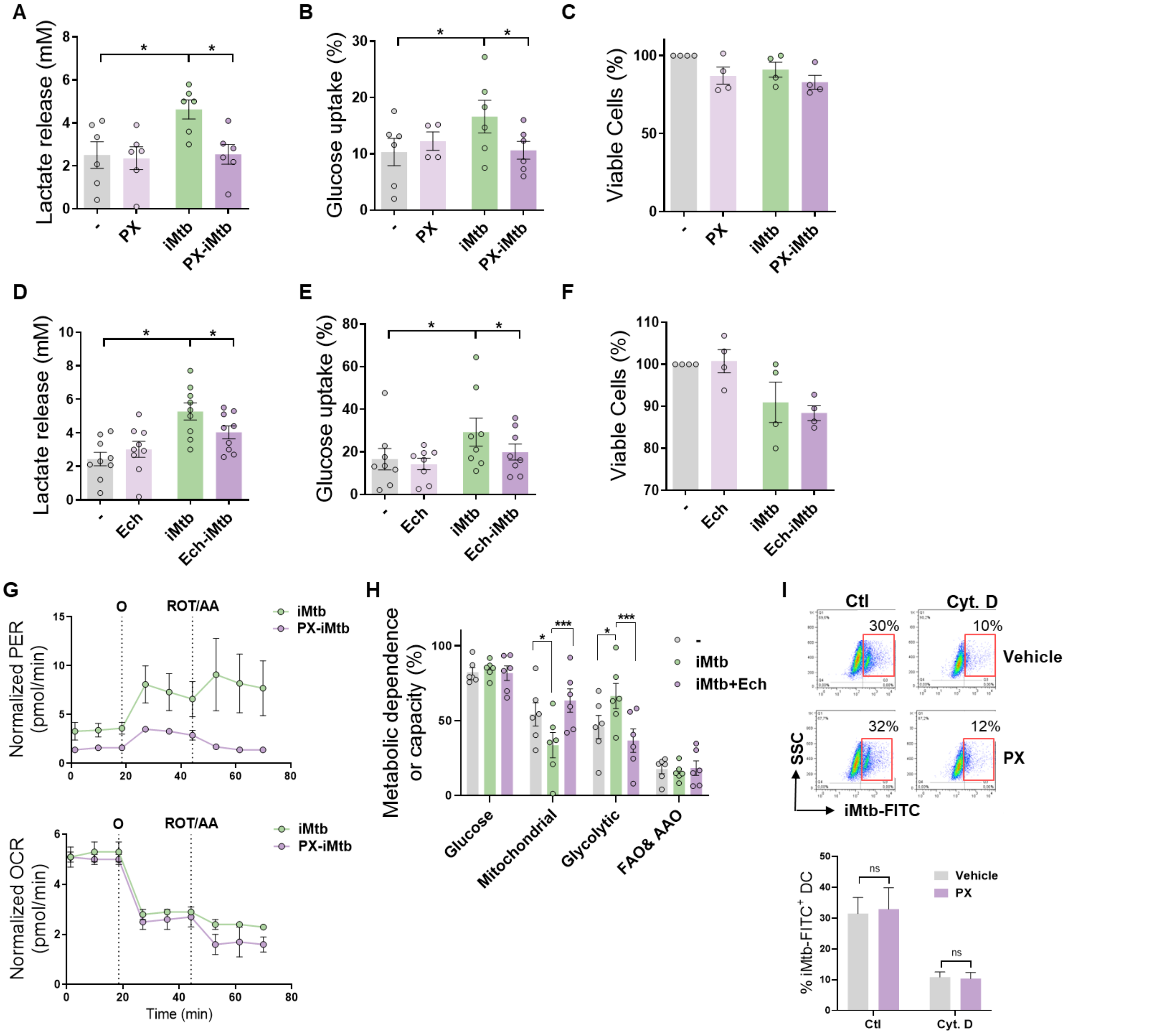


**HIF-1α activity is required to trigger the glycolytic pathway in iMtb-stimulated DCs.** Mo-DCs were stimulated with iMtb in the presence of PX-478 (PX, A-C) or Echinomycin (Ech, D-F), both HIF-1α inhibitors. **(A and D)** Lactate release measured in supernatant (N=6-8). **(B and E)** Glucose uptake measured in supernatant (N=6-8). **(C and F)** Percentage of viable cells relative to untreated DCs (N=4). **(G)** Kinetic profile of proton efflux rate (PER) and oxygen consumption rate (OCR) measurements in iMtb-stimulated DCs and PX-iMtb-stimulated DCs. **(H)** Relative contributions of glycolytic and FAO & AAO capacities and glucose and mitochondrial dependences to overall DC metabolism analyzed with SCENITH in DCs exposed to iMtb in the presence or not of Echinomycin (Ech) (N=4). **(I)** Uptake of Mtb-FITC by DCs treated or not with PX-478 in the presence or not of cytochalasin D (Cyt D), a potent phagocytosis inhibitor that interferes with actin polymerization. Representative dot plots and quantifications are shown (N=4). **(A-G, I)** 2way ANOVA followed by Tukey's multiple comparisons test (*$p < 0.05$), as depicted by lines. Values are expressed as means ± SEM. **(H)** Paired t test (*$p < 0.05$; ***$p < 0.001$) as depicted by lines. The data are represented as scatter plots with each circle representing a single individual, means ± SEM are shown.

**Fig. S4.**

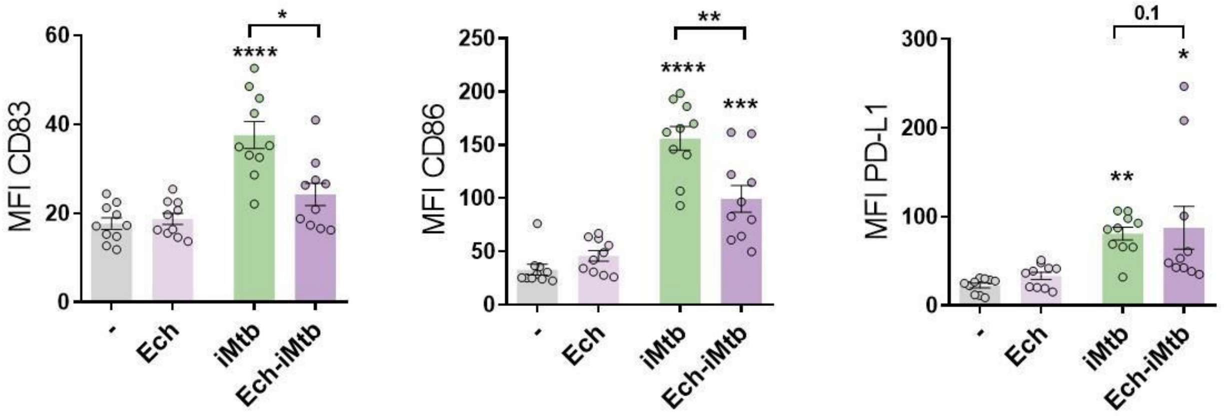


**HIF-1α is required to adopt a mature phenotype in iMtb-stimulated DCs.** Mo-DCs were stimulated with irradiated Mtb (iMtb) in the presence or not of Echinomycin (Ech), a HIF-1α inhibitor. Mean fluorescence intensity (MFI) of CD83, CD86 and PD-L1 (N=10). 2way ANOVA followed by Tukey's multiple comparisons test (*p < 0.05; **p < 0.01; ***p < 0.001; ****p < 0.0001), compared to control cells or as depicted by lines. The data are represented as scatter plots with each circle representing a single individual, means ± SEM are shown.

**Fig. S5.**

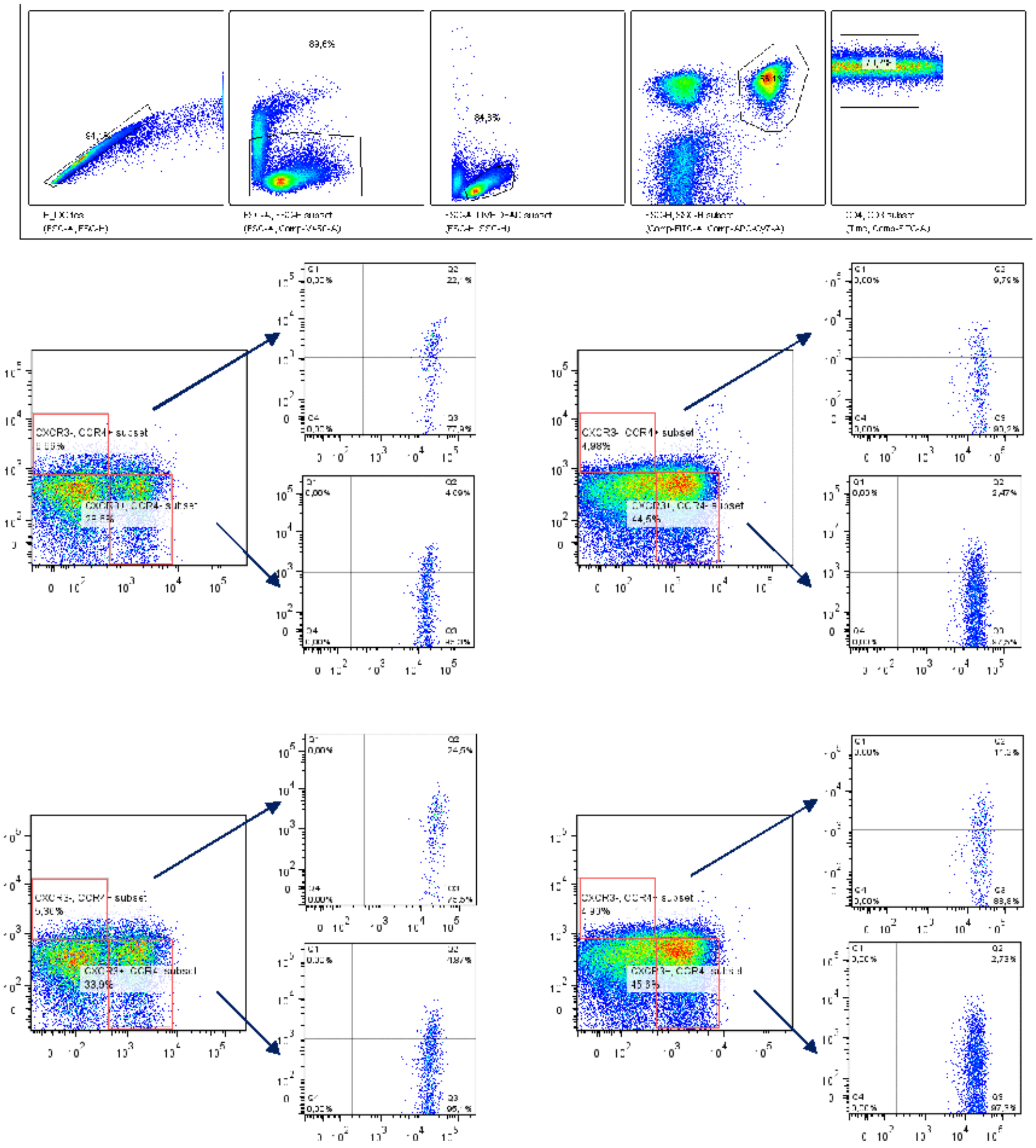

**Gating strategy to define CD4$^+$ T cells in response to Mtb.** Monocytes from PPD$^+$ healthy donors were differentiated towards DCs, challenged or not with iMtb in the presence or not of PX-478, and co-cultured with autologous CD4$^+$ T cells for 5 days. Gating strategy to define Th1 (CXCR3$^+$CCR4$^-$CCR6$^-$ cells), Th17 (CXCR3$^-$CCR4$^+$CCR6$^+$ cells), Th2 (CXCR3$^-$CCR4$^+$CCR6$^-$ cells) and Th1/Th17 (CXCR3$^+$CCR4$^-$CCR6$^+$ cells or Th1*) CD4$^+$ T populations by FACS.

**Fig. S6.**

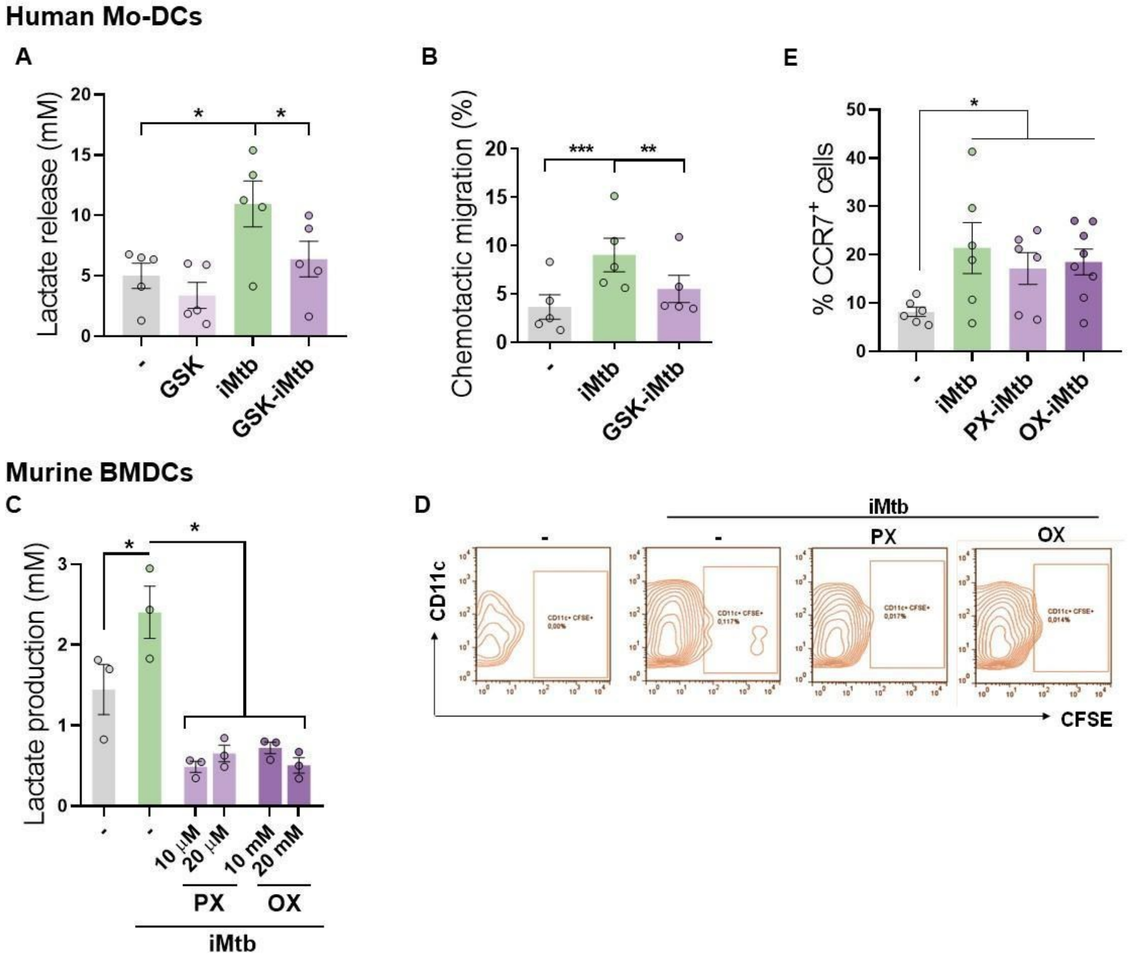


**Glycolysis is required to trigger the migratory activity in iMtb-stimulated DCs. (A-B)** Mo-DCs were treated or not with the glycolysis inhibitor GSK2837808A and stimulated with iMtb. **(A)** Lactate release (N=5). **(B)** Chemotactic activity towards CCL21 (N=5). **(C-D)** Murine BMDCs were treated or not with PX-478 or oxamate and stimulated with iMtb for 24 h. **(C)** Lactate release (N=3). **(D)** Representative dot blots showing the percentages of migrating BMDCs (CD11c$^{+}$, CFSE-labeled) determined from inguinal lymph nodes. **(E)** Mo-DCs were stimulated with iMtb in the presence of either PX-478 or oxamate and CCR7 expression was measured by FACS (N=6). 2way ANOVA followed by Tukey's multiple comparisons test ($*p < 0.05$; $**p < 0.01$; $***p < 0.001$), as depicted by lines. The data are represented as scatter plots with each circle representing a single individual, means ± SEM are shown.

**Fig. S7.**

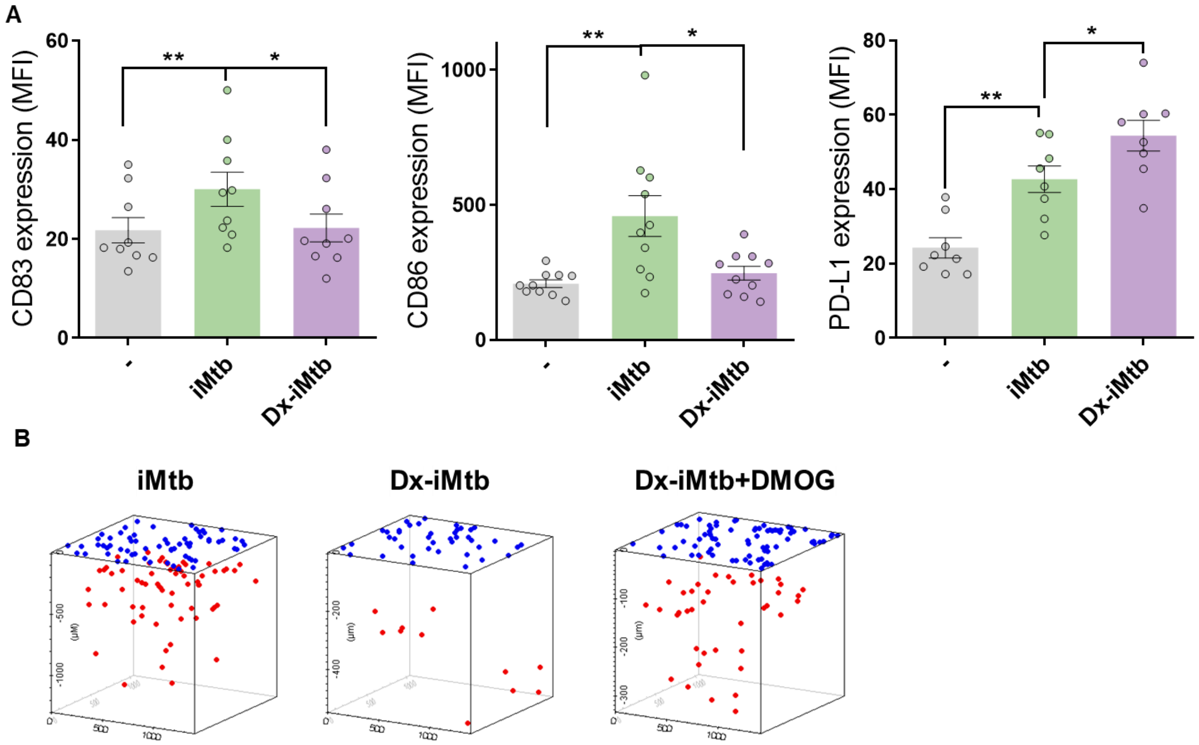


**Profile of tolerogenic DCs induced by Dexamethasone.** Tolerogenic Mo-DCs were generated in the presence or not of dexamethasone (Dx) and stimulated with iMtb. **(A)** Mean fluorescence intensity (MFI) of CD83, CD86 and PD-L1 (N=9). The data are represented as scatter plots with each circle representing a single individual. Values are expressed as means ± SEM **(B)** Representative schemes of migrating cells through a collagen matrix. 2way ANOVA followed by Tukey's multiple comparisons test ($*p < 0.05$; $**p < 0.01$), as depicted by lines.

**Fig. S8.**

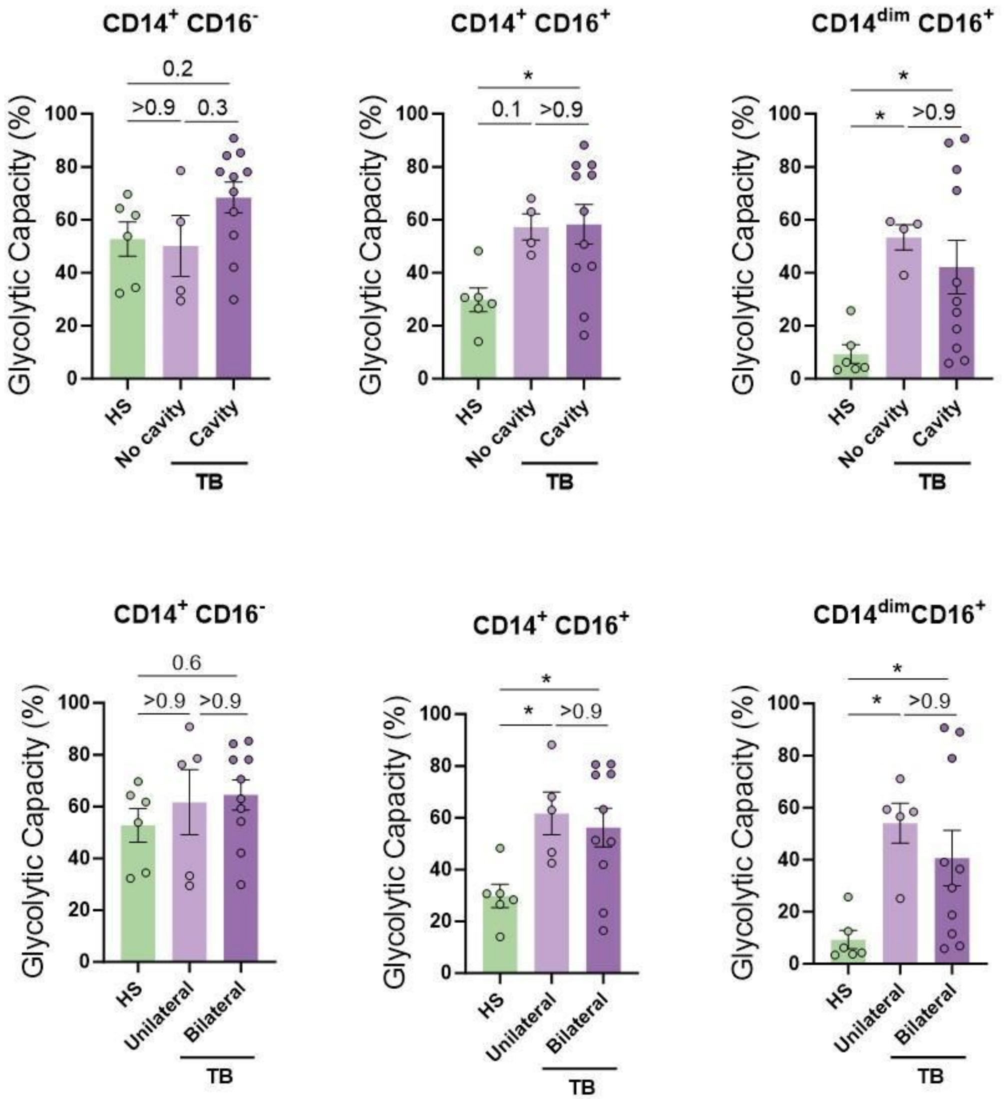


**Association between baseline glycolytic status of monocytes and the severity of lung disease**. The glycolytic capacity was assessed in monocyte subsets from TB patients with bilateral versus unilateral disease, reflecting the extent of disease (upper panels), or with cavitary versus non-cavitary disease, reflecting the disease severity (lower panels) (N=4-11). The data are represented as scatter plots with each circle representing a single individual. P values were calculated using the Kruskal Wallis test with Dunn's correction for multiple comparisons.

**Fig. S9.**

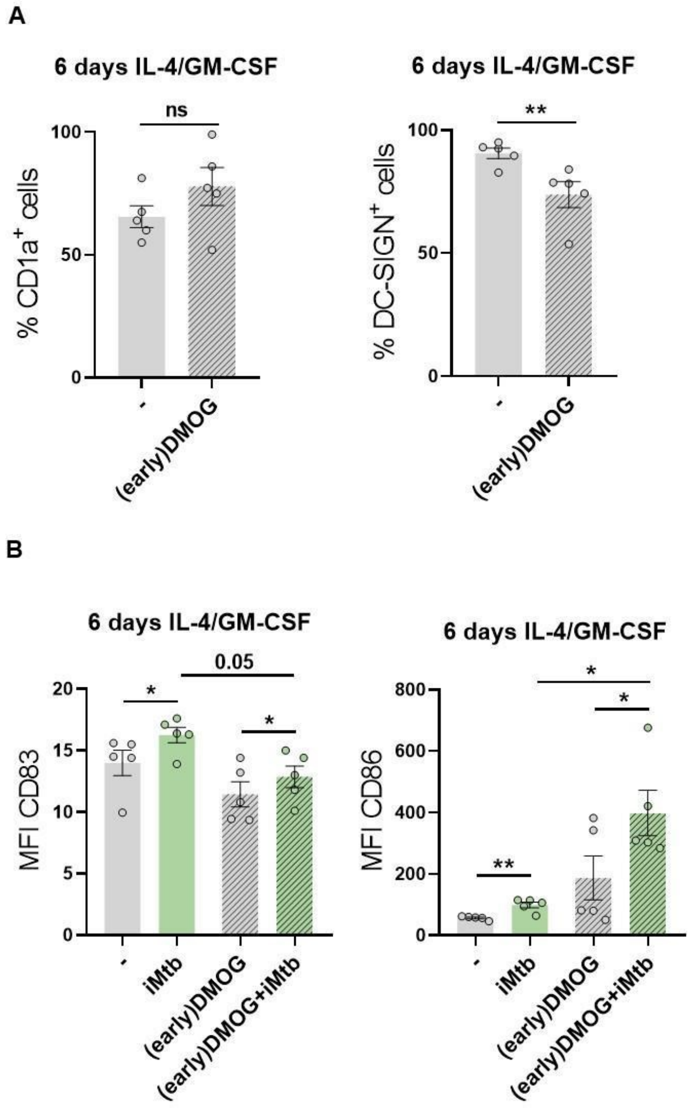


**Impact of premature activation of HIF-1α in monocytes on the generated DCs.** Monocytes from HS were treated with DMOG during the first 24h of differentiation with IL-4/GM-CSF (earlyDMOG) and removed afterwards. On day 6 of differentiation, cells were stimulated or not with iMtb. **(A)** Percentage of cells expressing CD1a and DC-SIGN after full 6 days of differentiation (N=5). **(B)** MFI of CD83 and CD86 on day 6 differentiated cells stimulated or not with iMtb for further 24h (N=5). The data are represented as scatter plots with each circle representing a single individual. Statistical significance was assessed by **(A)** paired T test ($**p < 0.01$); **(B)** 2-way ANOVA followed by Tukey's multiple comparisons test ($*p < 0.05$).
